# Characterizing the frequency-specific and spatiotemporal dynamics of β-γ Phase-Amplitude Coupling in Parkinson’s disease

**DOI:** 10.1101/2025.05.14.653744

**Authors:** Philipp A. Loehrer, Sahar Yassine, Immo Weber, Valentin Sanner, Shenghong He, Alek Pogosyan, Lijiao Chen, Laura Witt, Gereon Rudolf Fink, David J. Pedrosa, Lars Timmermann, Huiling Tan

## Abstract

Cross-frequency coupling (CFC) has been proposed to facilitate neural information transfer across spatial and temporal scales. Phase-amplitude coupling (PAC), a type of CFC in which the amplitude of a faster brain oscillation is coupled to the phase of a slower brain oscillation, is implicated in various higher-order cognitive functions and was shown to be pathologically altered in neurological and psychiatric disease. In Parkinson’s disease (PD), the coupling between gamma amplitude (50-150 Hz) to beta phase (13-35 Hz) is exaggerated. Enhanced β-γ PAC was found in the subthalamic nucleus and various cortical sources and shown to be responsive to dopaminergic therapy and deep brain stimulation (DBS). Therefore, exaggerated β-γ PAC has been proposed to be a disease marker and a potential target for brain circuit interventions. Despite these promising findings, a significant knowledge gap remains, as the spatial and frequency-specific dynamics of β-γ PAC and its association with motor symptoms and therapy remain elusive. To address this knowledge gap, we employed high-density electroencephalography (EEG) with advanced source localisation techniques for PD patients at rest. We highlight three key findings: (1) a frequency-specific increase in high β (23-35 Hz)-γ PAC within and between sources of the cortical motor network, (2) a link between elevated high β-γ PAC and bradykinesia and rigidity when OFF medication, but not tremor, and (3) a medication-induced reduction in high β-γ PAC in the supplementary motor area correlating with clinical improvement. Altogether, this study provides novel insights into the pathophysiology of PD as an oscillopathy and identifies high β-γ PAC as a potential marker of Parkinsonian symptoms and treatment effects. This has important implications for invasive as well as non-invasive therapeutic strategies as high β-γ PAC targeting might hold greater promise than targeting β-γ PAC per se.

## Introduction

Neural oscillations reflect the synchronised activity of neural ensembles and are supposed to facilitate information processing and transfer within and between distant regions of the human brain (Singer, 1999; Siegel *et al*., 2012). In this context, cross-frequency coupling (CFC). i.e. the interaction between neural oscillations of different frequencies, seems to be an important mechanism underlying information processing as larger neural populations typically oscillate at lower frequencies, while smaller ensembles are active at higher rhythms (Buzsaki, 2006; Aru *et al*., 2015). In particular, Phase-Amplitude Coupling (PAC), a type of CFC where the amplitude of a high-frequency oscillation is coupled to a specific phase of a low-frequency rhythm (Siems *et al*., 2016), is supposed to play a significant role in various higher cognitive functions. Most notably, coupling between theta phase (4-8 Hz) to gamma amplitude (30-100 Hz) in the human hippocampus has been implicated in memory and learning (Mormann *et al*., 2005; Tort *et al*., 2009; Lisman and Jensen, 2013; Igarashi *et al*., 2014; Daume *et al*., 2024). Furthermore, PAC is modulated in various activities, including visual (Spyropoulos *et al*., 2018) and auditory processing (Kikuchi *et al*., 2017), movement and speech (Nie *et al*., 2024), as well as complex cognitive functions (Voytek *et al*., 2015). Additionally, changes in PAC patterns have been linked to neurological and mental disorders, including Parkinson’s disease (PD), Alzheimer’s disease, schizophrenia, and obsessive compulsive disorder (for a review see: Salimpour and Anderson, 2019).

In PD, exaggerated coupling between beta phase (13-35 Hz) and gamma amplitude (50-150 Hz) has been detected in cortical and subcortical structures. In this regard, it is important to note that the subdivision of the beta frequency band into low-beta (13-22 Hz) and high-beta (23-35 Hz) is essential for Parkinson’s disease pathophysiology, as these sub-bands reflect distinct neurophysiological mechanisms with different clinical implications (van Wijk *et al*., 2016). Evidence demonstrates that high-beta activity in the STN predicts therapeutic responses to both levodopa and DBS, while low-beta in the STN correlates more strongly with overall motor symptom severity (van Wijk *et al*., 2016; Morelli and Summers, 2023). Despite this functional segregation, most investigations of β-γ PAC have analysed the beta band as a unified frequency band (13-35 Hz). Nonetheless, these studies revealed important patterns across brain regions. In particular, β-γ PAC was elevated in the subthalamic nucleus (STN) and positively associated with symptom severity of bradykinesia and rigidity (van Wijk *et al*., 2016). On the cortical level, exaggerated β-γ PAC was found in various cortical areas (Gong *et al*., 2021), including sensorimotor cortex and shown to be reduced by DBS (de Hemptinne *et al*., 2013; de Hemptinne *et al*., 2015). Furthermore, elevated β-γ PAC was detected on the scalp level in C3/C4 using EEG, associated with bradykinesia, and shown to be amenable to dopaminergic therapy (Swann *et al*., 2015; Miller *et al*., 2019). Given the associations between elevated PAC and symptom severity and the observed restoration to healthy levels with therapy, PAC is a promising marker for the disease itself and a potential target for neuromodulation therapies (Duchet and Bogacz, 2024). Despite these promising findings, a critical knowledge gap remains, as the spatial and frequency-specific dynamics of β-γ PAC on the cortical level, and its association with motor symptoms and therapy, remain elusive. This presents a substantial challenge to advancing new brain circuit interventions for PD. To address this knowledge gap, we combined high-density electroencephalography (EEG) with advanced source localisation techniques to characterize the spatial and frequency-specific patterns of exaggerated β-γ-phase-amplitude-coupling in PD. Using this approach, we provide evidence for a frequency-specific elevation of high β-band (23-35 Hz) to broadband-γ PAC within and between sources of the human motor network. This elevated high β to broadband-γ PAC is associated with bradykinesia and rigidity, and restoring healthy levels in the supplementary motor area (SMA) is associated with clinical improvement. Altogether, this study provides novel insights into the pathophysiology of PD as an oscillopathy and identifies high β-γ PAC as a potential marker of Parkinsonian symptoms and treatment effects.

## Methods

### Ethical Approval

The study was approved by the local ethics committee (study number: 14-130) and carried out following the Declaration of Helsinki.

### Participants

Seventeen patients with PD and fifteen healthy control subjects (HC) matched concerning age, sex, and cognitive function participated in this study upon written informed consent. Clinical diagnosis of PD was established according to recent diagnostic criteria (Postuma *et al*., 2016), and motor function was assessed before and during the experiment using the Unified Parkinson’s Disease Rating scale part III (UPDRS-III). All participants were right-handed, according to the Edinburgh Handedness Inventory. There was no indication of depressive symptoms or dementia according to Beck’s Depression Inventory-II (BDI-II) and Mini-Mental State Examination (MMSE). Participants were excluded if they had pathological MRI, a concomitant neurological or psychiatric disease, or impaired visual or auditory function. Two healthy controls and one patient had to be excluded due to severe artefacts interfering with the EEG recording. One patient had to be excluded as he discontinued MRI acquisition and subsequent EEG recordings due to claustrophobia. Following preprocessing, the datasets of 15 PD patients (mean age ± SD: 68.2 ± 8.2 years; 2 female) and 13 healthy controls (mean age ± SD: 64.4 ± 6.5 years; 1 female) were included for further analysis (for demographics cf. Table 1). PD patients and healthy controls were well matched for age and gender (p = .21).

**Table 1.**
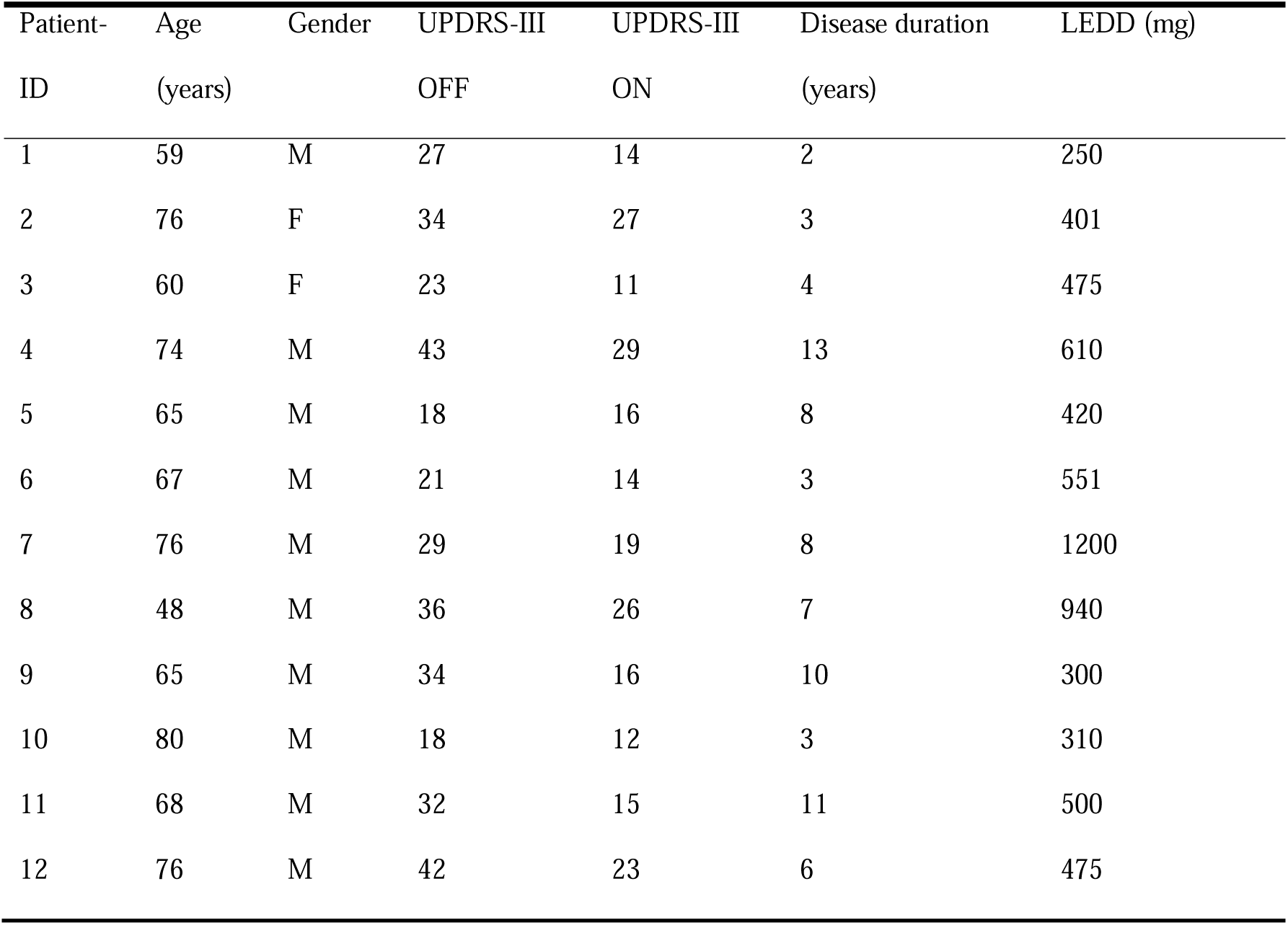
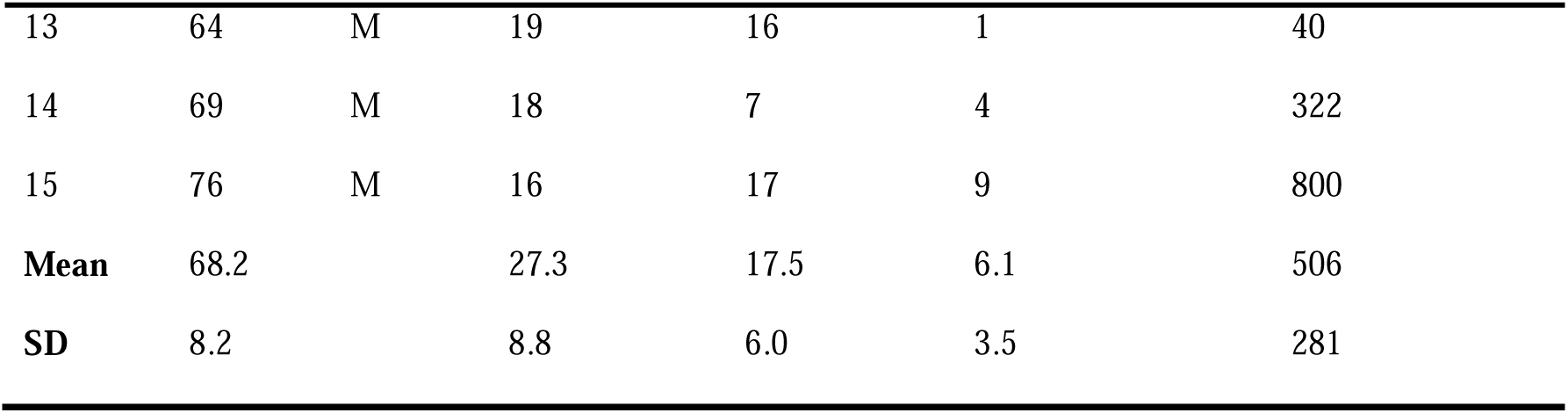
Sociodemographic information of patients, severity of parkinsonism and standard medication. Abbreviations: F = female; LEDD = Levodopa equivalent daily dose; M = male; UPDRS-III = Unified Parkinson’s Disease Rating Scale part III.

| Patient-ID | Age<br>(years) | Gender | UPDRS-III<br>OFF | UPDRS-III<br>ON | Disease duration<br>(years) | LEDD (mg) |
| --- | --- | --- | --- | --- | --- | --- |
| 1 | 59 | M | 27 | 14 | 2 | 250 |
| 2 | 76 | F | 34 | 27 | 3 | 401 |
| 3 | 60 | F | 23 | 11 | 4 | 475 |
| 4 | 74 | M | 43 | 29 | 13 | 610 |
| 5 | 65 | M | 18 | 16 | 8 | 420 |
| 6 | 67 | M | 21 | 14 | 3 | 551 |
| 7 | 76 | M | 29 | 19 | 8 | 1200 |
| 8 | 48 | M | 36 | 26 | 7 | 940 |
| 9 | 65 | M | 34 | 16 | 10 | 300 |
| 10 | 80 | M | 18 | 12 | 3 | 310 |
| 11 | 68 | M | 32 | 15 | 11 | 500 |
| 12 | 76 | M | 42 | 23 | 6 | 475 |
| 13 | 64 | M | 19 | 16 | 1 | 40 |
| 14 | 69 | M | 18 | 7 | 4 | 322 |
| 15 | 76 | M | 16 | 17 | 9 | 800 |
| **Mean** | 68.2 |  | 27.3 | 17.5 | 6.1 | 506 |
| **SD** | 8.2 |  | 8.8 | 6.0 | 3.5 | 281 |

### Data Acquisition

EEG-data were acquired using a 128-channel system with active electrodes (Brain Products GmbH, Gilching, Germany) and a reference channel on FCz, in an acoustically and electrically shielded room. Individual positions of electrodes and fiducial points were acquired using a 3D ultrasound digitisation system (Zebris Medical GmbH, Isny, Germany). After assuring that electrode impedances were below 10 kΩ, EEG-signals were recorded with Brainvision recorder (Brain Products GmbH, Gilching, Germany). Here, signals were amplified, band-pass filtered from 0.1 to 1000 Hz, and digitised at a sampling rate of 5000 Hz. The experimental paradigm consisted of multiple blocks with 15 seconds of continuous resting state and various movement tasks, whereby only data from the rest condition was analysed for this report. During the resting state recordings, participants were instructed to relax and focus on a cross presented on a screen before them. EEG of PD patients was recorded in medication OFF, after 12 hours of withdrawal of antiparkinsonian medication, and in medication ON, following the administration of 200 mg levodopa. In all patients, an experienced movement disorders specialist (PAL) assessed the UPDRS-III in the OFF and ON state.

Before EEG recordings, individual T1-weighted MRI of the head were acquired on a 3-Tesla Trio scanner (Siemens, Erlangen, Germany) using a 3D Modified Driven Equilibrium Fourier Transform sequence (repetition-time = 1930 ms, echo-time = 5.8 ms, flip-angle = 18°, slice-thickness = 1 mm) for individual source reconstruction.

### EEG Preprocessing

EEG data were preprocessed using the open-source Fieldtrip toolbox (Oostenveld *et al*., 2011, version: 20211020). First, the data was demeaned and detrended, high-pass filtered at 1 Hz, low-pass filtered at 500 Hz, and band-stop filtered at 50 Hz and its harmonics to remove residual power-line contamination. Then, data were downsampled to a sampling frequency of 1000 Hz and epoched into non-overlapping segments of 3000 ms, as it has been shown that PAC can be detected robustly in epochs of 2500 ms and longer, even at different levels of noise and coupling strength (Seymour *et al*., 2017; Hülsemann *et al*., 2019). Subsequently, an independent component analysis (ICA) identified and removed components containing eye movements, channel noise, muscle-, and electrocardiogram-artefacts. Segments were visually inspected, and channels and segments with poor data quality were excluded. Excluded channels were interpolated (20.5 +/- 1.9 channels for each patient and 19.4 +/- 2.9 channels for each healthy control), and a final visual inspection of the data was performed. If artefacts affected over 50% of the data segments, the entire dataset was excluded from subsequent analysis. As a result of this process, two healthy controls and one patient were excluded from the study. Following the removal of all artefacts, there were no significant differences in data volume, i.e. segments, between patients and controls (28.9 +/- 2.8 segments for each patient and 32.9 +/- 7.2 segments for each healthy control, p = 0.78). In a last step, data were re-referenced to the average reference of all electrodes.

### EEG-MRI Coregistration and 3D Cortical Mesh Construction

To reconstruct the dynamics of the brain at the cortical-source level, the inverse problem was solved based on the artefact-free EEG data and patients structural MRI employing functions from Freesurfer (http://surfer.nmr.mgh.harvard.edu/, version: 7.4.1), Brainstorm (version: 10-Oct-2023), and custom MATLAB (R2020b) scripts. The procedure is outlined in **Figure 1**. First, individual cortical surfaces were calculated from structural MRI scans using Freesurfer, creating a 3D cortical mesh with 15,000 vertices. The surface data was then registered to the Human Connectome Project (HCP) atlas, which is based on a multi-modal cortex parcellation and consists of 180 cortical areas per hemisphere (Glasser *et al*., 2016). As we were particularly interested in analysing β-γ PAC in the cortical motor network, we retained a fine resolution of the HCP-atlas in the regions “somatosensory and motor cortex” as well as “paracentral lobular and midcingulate cortex”. Still, we merged the remaining areas into broader regions as described by Glasser et al. This procedure yielded parcellations of 26 regions covering each hemisphere.

**Figure 1.**
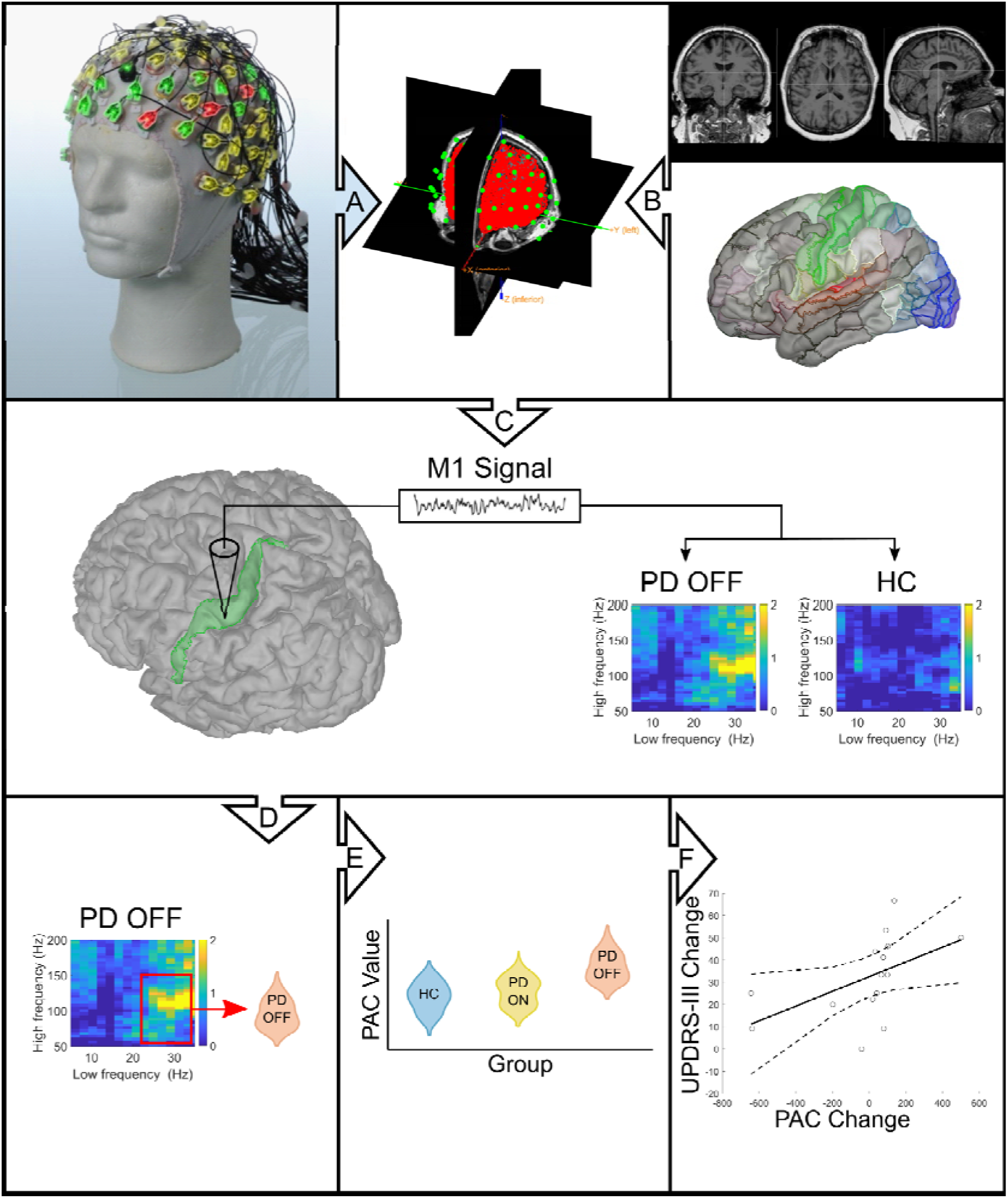
Schematic overview of data collection and analysis. **(A)** EEG data were recorded using a high-density EEG with active electrodes. **(B)** Individual structural MRI data were acquired. During MRI processing, the individual cortical surface was registered to the HCP-atlas to define regions of interest. **(C)** Artefact-free EEG and MRI data were co-registered by aligning the individual digitised electrode positions with the surface data to construct a realistic head model. An example of comparing PAC in M1 (green) between PD OFF and HC was shown here. **(D)** A linearly constrained minimum variance beamformer was used to extract source time series from a given ROI, and phase-amplitude coupling was examined using the Kullback-Leibler-based modulation index. Employing nonparametric permutation testing, we converted the ensuing modulation of index-values to a z-score by comparing them to a surrogate distribution. **(E)** PAC values were derived for each subject by averaging the comodulograms of a given ROI and group statistics **(F)** were performed using a repeated measures and a mixed-design ANOVA. The relationship between β-γ PAC and clinical features and treatment response were examined using Spearman correlations. **Abbreviations:** EEG = electroencephalography; HC = healthy controls; HCP = human connectome project; OFF = Parkinson’s disease patients in the OFF-medication state; PD ON = Parkinson’s disease patients in the ON-medication state; PAC = phase-amplitude coupling; ROI = region of interest; ANOVA = analysis of variance.

EEG data were then co-registered with the segmented MRI by aligning the individual digitised electrode positions with the surface data obtained from the structural scans. Subsequently, a realistic head model was constructed from the MRI-EEG aligned data employing an OpenMEEG boundary element method (Gramfort *et al*., 2010).

### Region of interest-based Source Reconstruction

Source analysis was performed utilising a linearly constrained minimum variance beamformer (LCMV; (Veen *et al*., 1997)). This technique applies a spatial filter to the EEG data at each vertex of the 3D cortical mesh, aiming to maximise the signal from that particular location while suppressing signals from other areas (Seymour *et al*., 2017). The spatial filter was constructed by combining the covariance matrix obtained from the sensor data with the leadfield information obtained from the head model. To extract a single source signal for each region of interest in the cortical motor network (dorsolateral prefrontal cortex (DLPFC), lateral premotor cortex (lPM), primary motor cortex (M1), supplementary motor area (SMA)), we constructed a single spatial filter for each ROI by calculating the mean of the concatenated filters from all vertices identified to lie within a ROI. Sensor-level data was then multiplied with this spatial filter to extract the source time series for each ROI.

### Phase-Amplitude Coupling Analysis

Source time series were examined for β-γ phase-amplitude coupling employing the Kullback-Leibler-based modulation index (Tort *et al*., 2008). As a first step, trials were zero-padded with segments of 750 ms, which were discarded after filtering, as spurious PAC can occur through filtering sharp edge artefacts (Kramer *et al*., 2008). Subsequently, low-frequency phase and high-frequency amplitude were extracted from each trial’s source time-series, employing a 2-way finite impulse response filter (eegfilt.m with fir1 parameters). Here, source time series were filtered across the 4 to 50 Hz range using a sliding window with a step size of 2 Hz and a bandwidth of 2 Hz to extract phase. Similarly, source time series were filtered across the 4 to 200 Hz range using a sliding window with a step size of 4 Hz to extract amplitude. As the bandwidth for filtering the amplitude-providing time series should depend on the frequency of the phase-providing frequency (Seymour *et al*., 2017), the bandwidth was chosen to be wide enough to capture the centre frequency ± the modulating low-frequency phase. Subsequently, instantaneous phase and amplitude were extracted from the filtered signals using a Hilbert transform.

To calculate the modulation index, phases from -180° to 180° were first binned into 18 bins of 20° each (Tort *et al*., 2008). Subsequently, the mean amplitude of the amplitude-providing frequency in each phase bin of the phase-providing frequency was calculated and normalised by the sum of the mean amplitudes for all bins. This created an amplitude-phase distribution similar to a probability distribution, which was then quantified using Shannon entropy. As Shannon entropy represents a variable’s inherent amount of information, it is maximal, if – in this context – the amplitude in each phase bin is equal (Hülsemann *et al*., 2019). This would represent a uniform distribution and correspond to the absence of phase-amplitude coupling. To measure the disparity between the calculated distribution and a uniform distribution, the Kullback-Leibler distance is calculated to derive the modulation index (MI). To quantify the meaningfulness of the derived MI-values, remove forms of spurious coupling, and make MI-values amenable to statistical evaluation, we performed nonparametric permutation testing (Swann *et al*., 2015; Hülsemann *et al*., 2019). Here, we converted MI-values to a z-score by comparing the observed MI-value to a surrogate distribution of shuffled MI-values. This surrogate distribution was generated by temporally shifting the phase time series compared to the amplitude time series. The phase time series was circularly shifted by cutting it at a randomly selected point within its length, effectively introducing a random temporal offset. Phase-amplitude coupling was then calculated, and this procedure was repeated 200 times.

### Power spectral density

To exclude the possibility that differences in spectral power in either β- or γ-band drive enhanced PAC, we calculated the power spectral density (PSD) of the different re-constructed source regions using the Welch method. Here, the Hamming window was set to 1000 ms with 50 % overlap. The PSD values for each ROI and each frequency band were then normalised by dividing it by the average power over the whole frequency range (1-300 Hz, excluding 50 Hz and its harmonics) and compared between groups using Wilcoxon-signed rank tests and Mann-Whitney-U tests, when applicable, corrected for multiple comparisons using the Benjamini-Hochberg method (Benjamini and Hochberg, 1995).

### Non-Sinusoidal Oscillations

Another significant challenge encountered in the analysis of PAC involves the presence of non-sinusoidal oscillations, which can lead to spurious PAC (Jensen *et al*., 2016; Lozano-Soldevilla *et al*., 2016; Cole *et al*., 2017). These characteristics of an oscillation can be assessed by determining the time intervals from trough to peak (rise-time), peak to trough (decay-time), and the ratio between these intervals (Dvorak and Fenton, 2014; Cole and Voytek, 2017). Furthermore, the sharpness of an oscillation can be assessed by evaluating the ratio between peak and trough sharpness. Peak sharpness was determined by calculating the average voltage difference between the peak and the five data points (i.e. 5 ms) preceding and following it. Trough sharpness was measured using a similar approach (Cole and Voytek, 2017). We computed sharpness and steepness ratios for the time series data of each ROI in the 13 to 35 Hz range. Subsequently, we conducted Wilcoxon-signed rank tests and Mann-Whitney-U tests, when applicable, corrected for multiple comparisons using the Benjamini-Hochberg method (Benjamini and Hochberg, 1995), to assess differences in non-sinusoidal oscillations between groups.

### Simulated PAC Analysis

To assess the reliability of our PAC algorithm, we generated 3 seconds of simulated data containing predetermined β-γ PAC (with centre frequencies of fphase = 21 Hz and famplitude = 100 Hz; code adjusted from (Hülsemann *et al*., 2019)). Subsequently, we introduced a variable level of random noise and varied the modulation strength. Comodulograms were then generated using our PAC algorithm across 30 trials of simulated data.

### Statistical Analysis

Group statistics were performed to compare β-γ PAC among PD patients in the medication ON and OFF state, and between PD patients and healthy controls. A repeated measures ANOVA was employed to analyse the PAC differences within PD patients, with the repeated measures factors ‘source’ (e.g., cortical regions of interest) and ‘frequency’ (e.g., low (13-22 Hz) and high (23-35 Hz) β-γ PAC-values), the independent variable ‘medication state’ (ON or OFF), and the covariate ‘hemisphere’ (left or right), using Greenhouse-Geisser correction for non-sphericity when appropriate. A mixed design ANOVA was performed for comparisons between PD patients and HC, with the within-subjects factors ‘source’ and ‘frequency’, the between-subjects factor ‘group’ (PD or HC), and the covariate ‘hemisphere’ (left or right). All statistical analyses were conducted to test for main effects and interactions between factors. Post-hoc analyses were performed where appropriate to identify significant pairwise comparisons. For statistical comparison, a PAC value was derived for each subject by averaging the co-modulograms of the regions of interest. To delineate differences in high and low β-γ PAC, we averaged MI values over the low beta-band (13-22 Hz) as well as the high-beta band (23-35 Hz) for the phase frequency and the broadband gamma range (50-150 Hz) for the amplitude-frequency. These frequency ranges were chosen based on prior research (Swann *et al*., 2015; Gong *et al*., 2021; Chikermane *et al*., 2024), whereby power analyses and analyses of sinusoidal oscillations focused on the same frequency ranges. Averaged MI values were squareroot-transformed to fulfil assumptions of ANOVA testing. Spearman correlations were used to investigate the relationship between β-γ PAC and clinical features and treatment response, corrected for multiple comparisons using the Benjamini-Hochberg method (Benjamini and Hochberg, 1995).

### Data and code availability

The data supporting the present findings are available on request from the corresponding author (PAL). The data are not publicly available due to privacy or ethical restrictions. All tools used to analyse MRI data are based on FreeSurfer Version 7.1 (Fischl, 2012), which is freely available. Tools employed to analyse EEG data are based on custom MATLAB scripts as well as Brainstorm version 20231010 (Tadel *et al*., 2011), FieldTrip version 20211020 (Oostenveld *et al*., 2011), and OpenMEEG (Gramfort *et al*., 2010), which are available online.

## Results

### Phase-amplitude coupling within sources of the human motor network is enhanced in Parkinson’s disease

We assessed the effects of participant group (PD OFF vs. HC), frequency for phase (low β vs. high β), and each considered source on β-γ PAC using a three-way mixed-design ANOVA. This analysis demonstrated significant main effects of ‘group’ (F(1,26) = 5.76, p = .02, η_p_^2^ = .098, c.f. **Supplementary Figure 1**) and ‘frequency’ (F(1,27) = 5.39, p = .024, η_p_^2^= .092), but not ‘source’ (F(3,81) = .95, p = .41, η_p_^2^ = .018). Here, β-γ PAC was higher in PD patients OFF medication than HC. The effect, however, was present for high β-γ PAC (high β: 23-35 Hz; frequency x group interaction: F(1,26) = 2.72, p = .014, η_p_^2^= .11), but not for low β-γ PAC (low β: 13-22 Hz). Post hoc t-tests for the different ROIs demonstrated that high β-γ PAC was increased in PD patients in M1 (t(26) = -2.28, p = .014, Cohen’s d = .58) and lPM (t(26) = -2.96, p = .003, Cohen’s d = .75, c.f. **Figure 2**). The mixed design ANOVA comparing PD patients ON medication and HC revealed no significant difference in overall β-γ PAC between the two groups (F(1,26) = 0.05, p = .944, η_p_^2^ = <.001) and between sources (F(3,81) = 2.49, p = .079, η_p_^2^ = .045). However, a significant main effect of ‘frequency’ (F(1,27) = 9.78, p = .003, η_p_^2^ = .156) indicated differences in β-γ PAC across frequency bands. Additionally, no interaction between frequency-band and group existed (F(1,26) = 0.02, p = .961, η_p_^2^ = <.001).

**Figure 2.**
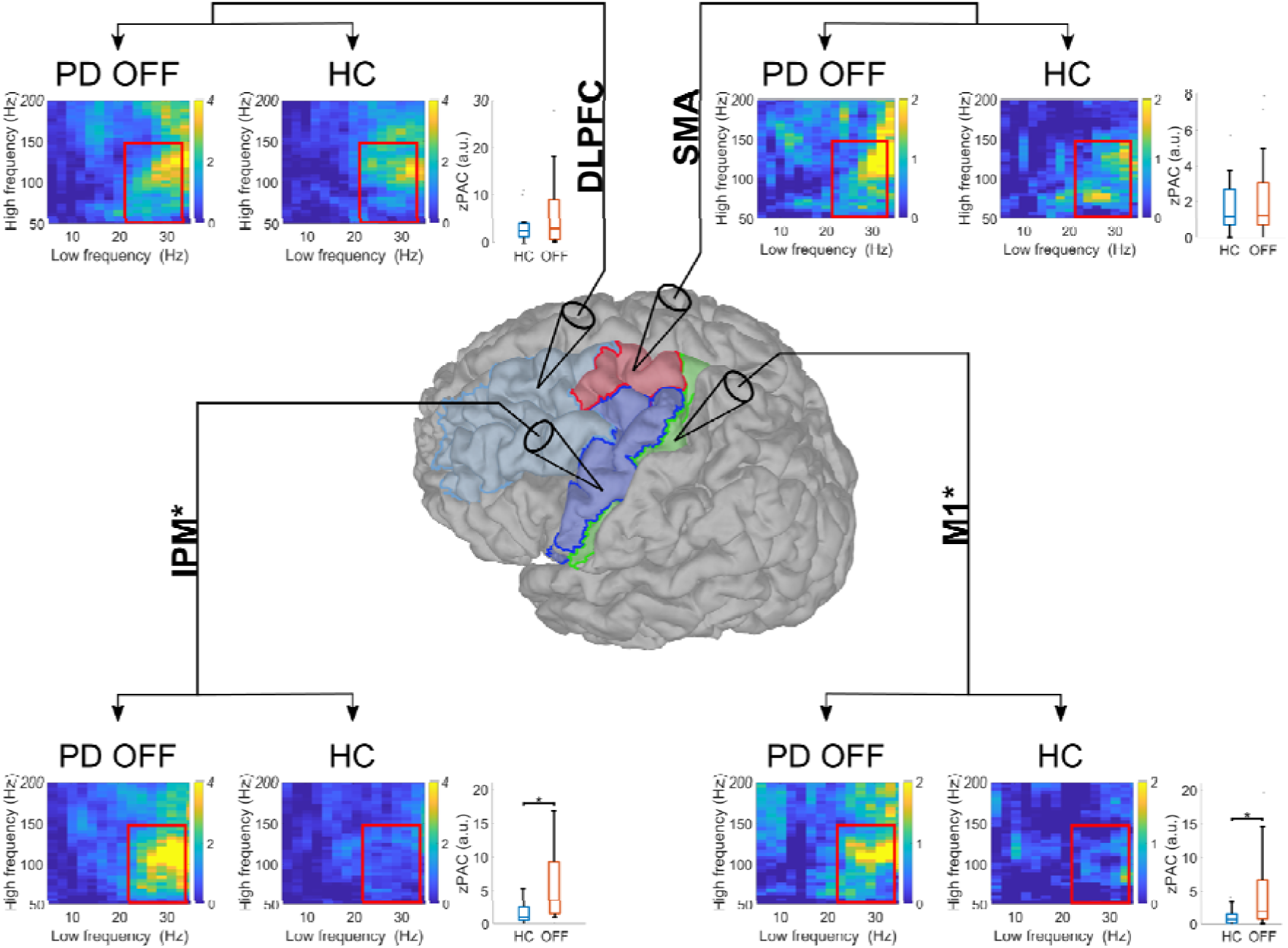
Comparison of comodulograms between PD patients OFF medication and healthy controls. Group comodulograms displaying the modulation index (MI) across subjects for each group within the four regions of interest. The comodulograms represent MIz-values, obtained through nonparametric permutation testing by comparing the observed MI-values against a surrogate distribution of shuffled MI-values. Brain regions with significant group differences, as assessed by post-hoc testing, are marked by an asterisk. For boxplots, zPAC values were averaged across hemispheres. The boxes represent the interquartile range (IQR), with the horizontal line indicating the median. Whiskers extend to 1.5 times the IQR or the most extreme data points within this range, while outliers are shown as individual points. **Abbreviations:** DLPFC = dorsolateral prefrontal cortex; HC = healthy controls; lPM = lateral premotor cortex; M1 = primary motor cortex; OFF = Parkinson’s disease patients in the OFF-medication state; PD ON = Parkinson’s disease patients in the ON-medication state; SMA = supplementary motor area

We assessed the effects of medication status (PD OFF vs. PD ON), frequency for phase (low β vs. high β) and each considered source on β-γ PAC using a three-way repeated measures ANOVA. The analysis demonstrated significant main effects of ‘medication’ (F(1,26) = 6.37, p = .014, η_p_^2^ = .101) and ‘frequency’ (F(1,27) = 5.66, p = .021, η_p_^2^ = .09), whereas no significant main effect of ‘source’ was observed (F(3,81) = .48, p = .683, η_p_^2^ = .008). Here, PD patients OFF medication had higher overall β-γ PAC values than ON medication. This effect was present for high β-γ PAC only (frequency x medication interaction: F(1,26) = 7.98, p = .007, η_p_^2^ = .123). Post hoc paired t-tests for the different ROIs demonstrated that high β-γ PAC was increased in PD patients OFF medication in M1 (t(14) = -1.81, p = .04, Cohen’s d = .47), lPM (t(14) = -2.39, p = .011, Cohen’s d = .6), and SMA (t(14) = -2.73, p = .005, Cohen’s d = .66), whereby a trend towards significance was observed for DLPFC (t(14) = -1.68, p = .052, Cohen’s d = .36; c.f. **Figure 3**). In the OFF medication state, correlation analysis revealed a significant relationship between high β-γ PAC and UPDRS-OFF values in M1 (p = .04, rho = .535), lPM (p = .04, rho = .569), and SMA (p = .049, rho = .475, c.f. **Supplementary Figure 2**). Furthermore, high β-γ PAC-values were correlated with the bradykinesia-rigidity subscore in M1 (p = .035, rho = .521), lPM (p = .035, rho = .509), and SMA (p = .035, rho = .543, c.f. **Supplementary Figure 3**) but not with the tremor subscore (all p > .43). There were no correlations between high β-γ PAC and UPDRS-ON values (all p > .066).

**Figure 3.**
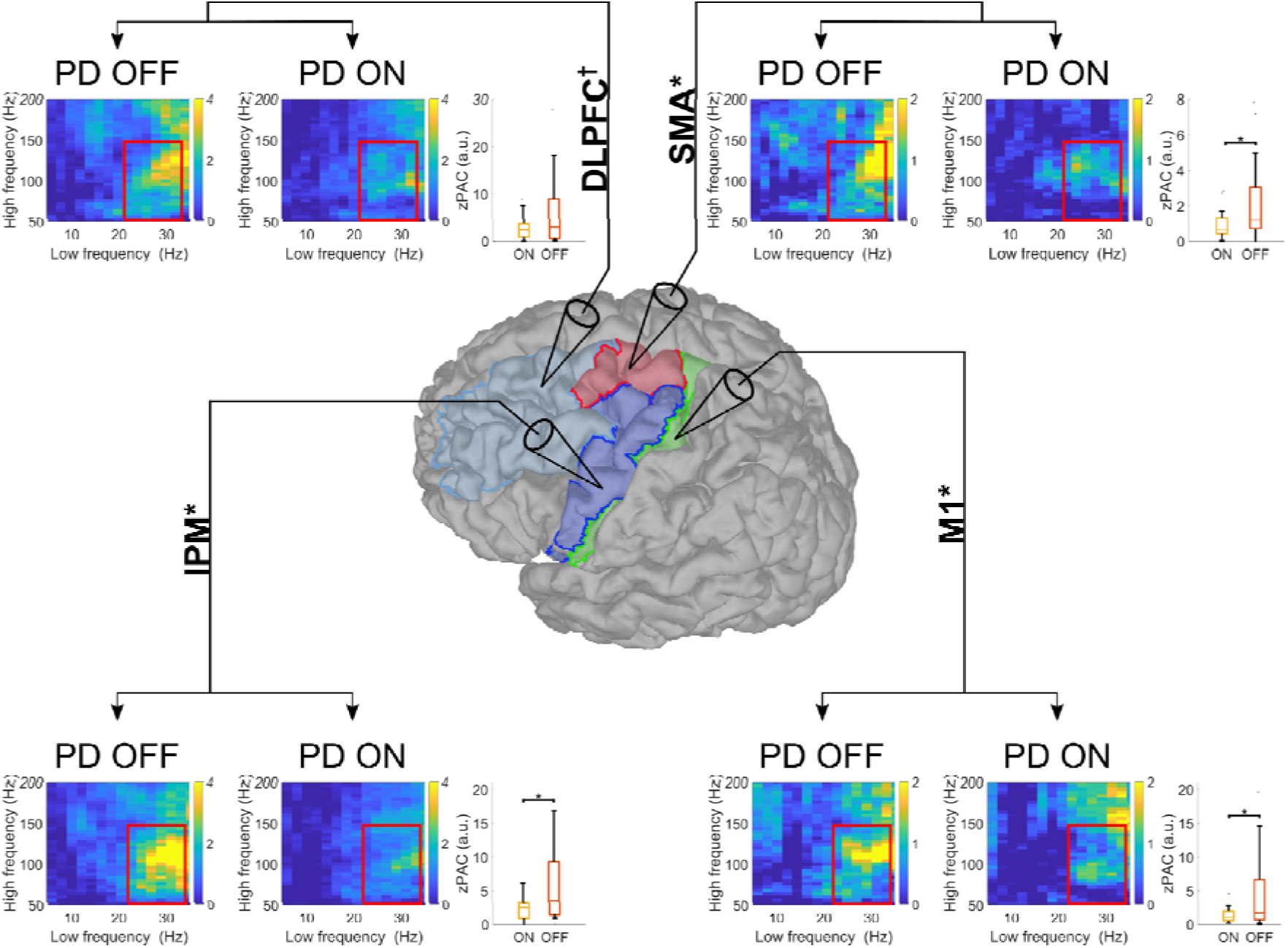
Comparison of comodulograms between PD patients OFF medication and PD patients ON medication. Group comodulograms displaying the modulation index (MI) across subjects for each group within the four regions of interest. The comodulograms represent MIz-values, obtained through nonparametric permutation testing by comparing the observed MI-values against a surrogate distribution of shuffled MI-values. Brain regions with significant group differences, as assessed by post-hoc testing, are marked by an asterisk. Please note that there was a trend towards significance for DLPFC (p = .052; marked by †). For boxplots, zPAC values were averaged across hemispheres. The boxes represent the interquartile range (IQR), with the horizontal line indicating the median. Whiskers extend to 1.5 times the IQR or the most extreme data points within this range, while outliers are shown as individual points. **Abbreviations:** DLPFC = dorsolateral prefrontal cortex; HC = healthy controls; lPM = lateral premotor cortex; M1 = primary motor cortex; OFF = Parkinson’s disease patients in the OFF-medication state; PD ON = Parkinson’s disease patients in the ON-medication state; SMA = supplementary motor area

As prior studies have demonstrated an association between medication effects on electrophysiological markers and clinical subscores of the predominantly affected side (Miller *et al*., 2019), we sought to evaluate the impact of medication on high β-γ PAC and clinical parameters. Specifically, we correlated the percentage change in high β-γ PAC of the ROIs contralateral to the primarily affected side with the percentage change in the UPDRS-III score and its subscores of the primarily affected side. Here, medication-induced change in high β-γ PAC of the SMA correlated with overall symptom improvement (p = .004, rho = .731) as well as improvement of bradykinesia and rigidity (p = .002, rho = .758) but not with tremor improvement (p = .11, rho = .569, c.f. **Figure 4**). This effect remained significant even after excluding outliers (UPDRS-III: p = .041, rho = .683; bradykinesia-rigidity subscore: p = .046, rho = .691, c.f. **Supplementary Figure 4**). All p-values are corrected for multiple comparisons using the Benjamini-Hochberg method.

**Figure 4.**
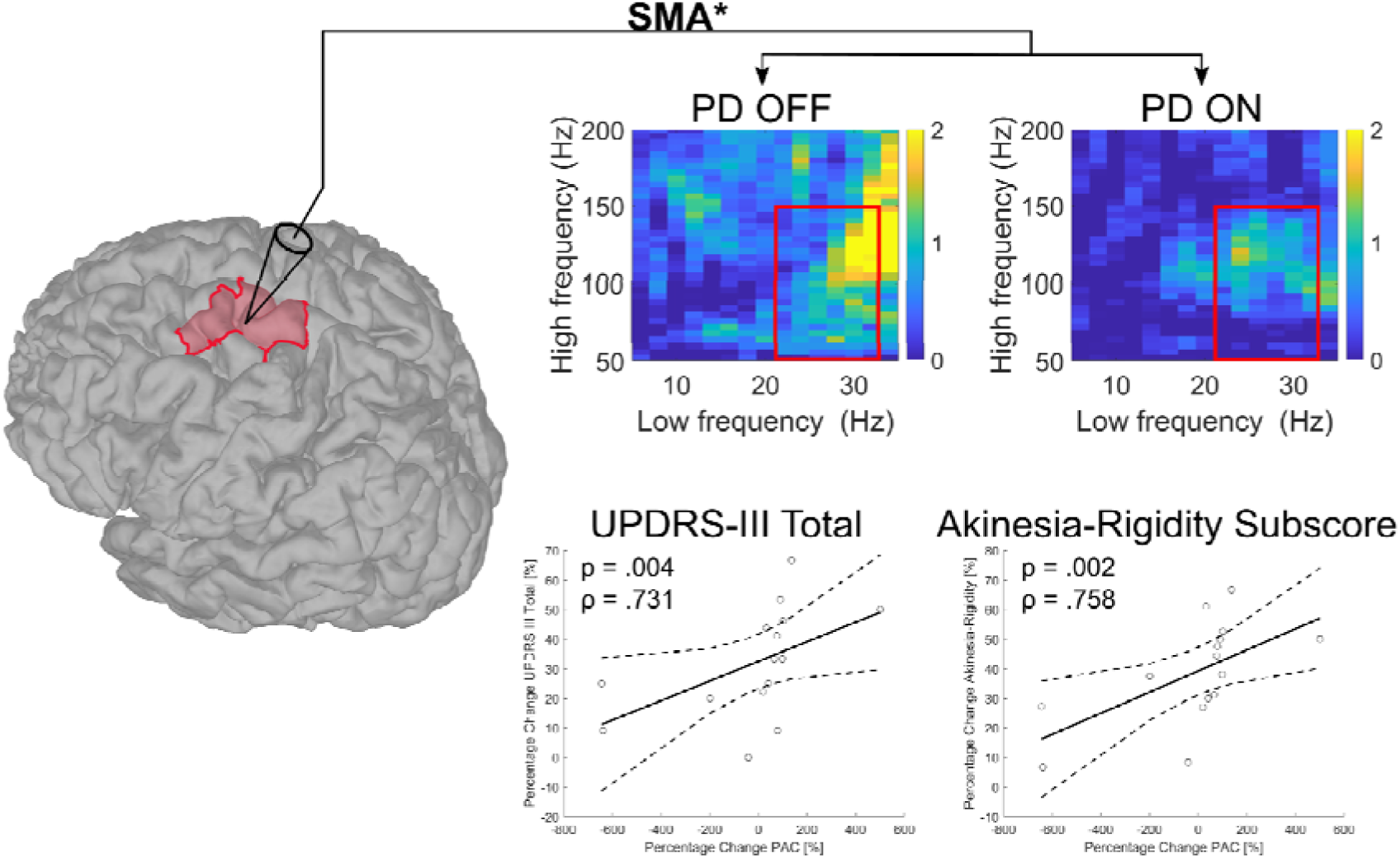
Correlation of high β-γ PAC changes in the SMA with UPDRS-III improvement. Relationship between the percentage change in high β-γ PAC within the SMA of the primarily affected side and the percentage change in UPDRS-III total scores and the bradykinesia-rigidity subscore. A greater reduction of high β-γ PAC in SMA was associated with a greater reduction in UPDRS-III total scores and bradykinesia-rigidity subscores. The solid black line represents the best-fit line while dashed lines represent 95% confidence intervals. P-values are corrected for multiple comparisons using the Benjamini-Hochberg method. **Abbreviations:** PAC = phase-amplitude coupling; OFF = Parkinson’s disease patients in the OFF-medication state; PD ON = Parkinson’s disease patients in the ON-medication state; SMA = supplementary motor area; UPDRS = Unified Parkinson’s Disease Rating Scale

### Phase-amplitude coupling between sources of the human motor network is enhanced in Parkinson’s disease

Phase-amplitude coupling between sources indicates connectivity between different sources in the human motor network. Differences in inter-source β-γ PAC between patients and HC were assessed using a three-way mixed design ANOVA with group (PD OFF vs. HC), interconnected sources, and frequency for phase (high β vs. low β) as factors. This analysis demonstrated no differences for the main effects ‘frequency’ (F(1,27) = 1.94, p = .169, η_p_^2^ = .035) and ‘source’ (F(11,297) = 1.19, p = .315, η_p_^2^ = .022). However, a significant main effect of ‘group’ (F(1,26) = 4.92, p = .031, η_p_^2^ = .085, c.f. **Supplementary Figure 5**) was observed. Here, inter-source β-γ PAC was higher in PD patients OFF medication than HC. The effect, however, was present for high β-γ PAC (frequency x group interaction: F(1,26) = 5.08, p = .028, η_p_^2^ = .087), but not for low β-γ PAC. Post hoc t-tests demonstrated that high β-γ PAC was increased in PD patients between several sources of the cortical motor network. **Table 2** summarises the significant results of post hoc t-testing. The mixed design ANOVA comparing PD patients ON medication and HC revealed no significant difference in inter-source β-γ PAC for the main factors ‘group’ (F(1,52) = .12, p = .727, η_p_^2^ = .002) and ‘source’ (F(11,297) = 1.28, p = .267, η_p_^2^ = .024). However, there was a significant main effect of ‘frequency’ (F(1,27) = 10.01, p = .003, η_p_^2^ = .159), indicating differences in β-γ PAC across frequency bands. Additionally, no interaction between frequency-band and group existed (F(1,26) = <.001, p = .994, η_p_^2^ = <.001).

**Table 2.** Post-hoc comparison of high β-γ PAC between groups. t-tests were employed to compare healthy controls (HC) and PD patients OFF medication, while paired t-tests were used to compare between PD patients ON and OFF medication. P-values were corrected for multiple comparisons using the Benjamini-Hochberg method. Bold font indicates significant p-values after adjustment for multiple comparisons.

| HC vs PD-OFF |  |  | PD-ON vs PD-OFF |  |  |
| --- | --- | --- | --- | --- | --- |
| *connection* | *p* | *effect size* | *connection* | *p* | *effect size* |
| **M1-IPM** | .042 | 0.66 | **M1-DLPFC** | .008 | 0.554 |
| **IPM-M1** | .042 | 0.62 | **DLPFC-M1** | .018 | 0.675 |
| **DLPFC-M1** | .042 | 0.58 | **M1-IPM** | .018 | 0.555 |
| **DLPFC-IPM** | .042 | 0.553 | **SMA-DLPFC** | .018 | 0.545 |
| **M1-DLPFC** | .042 | 0.562 | **DLPFC-SMA** | .021 | 0.529 |
| IPM-DLPFC | .055 | 0.502 | **IPM-M1** | .025 | 0.602 |
|  |  |  | **DLPFC-IPM** | .036 | 0.494 |
|  |  |  | IPM-DLPFC | .052 | 0.402 |
|  |  |  | M1-SMA | .056 | 0.356 |
|  |  |  | SMA-IPM | .057 | 0.349 |

We assessed the effects of medication status (PD OFF vs. PD ON), frequency for phase (low β vs. high β) and inter-connected sources on β-γ PAC using a three-way repeated measures ANOVA. There were no significant differences for the main effects of ‘frequency’ (F(1,27) = 2.47, p = .122, η_p_^2^ = .042) and ‘source’ (F(11,297) = .84, p = .497, η_p_^2^ = .015). The analysis, however, demonstrated a significant main effect of ‘medication’ (F(1,26) = 4.87, p = .031, η_p_^2^ = .079). Here, PD patients OFF medication had higher overall β-γ PAC values than ON medication. This effect was present for high β-γ PAC only (frequency x medication interaction: F(1,52) = 6.16, p = .016, η_p_^2^ = .098). Post hoc t-testing demonstrated that high β-γ PAC was increased in PD patients OFF medication between several sources of the cortical motor network, and significant results are summarised in **Table 2**. There were no correlations between interregional β-γ PAC and clinical scores after correcting for multiple comparisons.

### Power spectral density

We calculated the power spectral density using the Welch method to assess the possibility that enhanced β-γ PAC was driven by altered spectral power in β- or γ-band in PD patients. No differences in β- or γ-band in PD patients ON and OFF medication compared to HC were detected (all p > .09; c.f. **Supplementary Figure 6** and **Supplementary Figure 7**).

### Non-sinusoidal Oscillations

The sharpness and steepness ratios were calculated to determine whether PAC results were affected by deviations from perfect sinusoidal waveform of oscillations. No differences in the β-band between groups were detected in these ratios (all p > .57), suggesting that our results are unlikely to be undermined by differences in waveform shapes (c.f. **Supplementary Figure 8** and **Supplementary Figure 9**).

### Simulated phase-amplitude coupling

We further validated our PAC results by generating simulated data segments of three seconds, including known beta-gamma coupling (21 Hz phase, 100 Hz amplitude centre frequencies) at various levels of random noise and different coupling strengths. Here, we used the identical MATLAB code previously employed to analyse the EEG data and successfully detected beta-gamma PAC (c.f. **Supplementary Figure 10**).

## Discussion

In the present study, we used high-density EEG combined with advanced source localisation techniques to characterise the spatial and frequency-specific patterns of exaggerated β-γ phase-amplitude coupling in Parkinson’s disease. There are three key findings. First, we demonstrate a frequency-specific elevation of β-γ PAC within and between sources of the human motor network with the phase of high β-band but not low β-band oscillation modulating the power of γ band activities. Second, we identify an association of elevated high β-γ PAC with bradykinesia and rigidity within distinct sources of the motor network when patients were OFF medication. Tremor scores, however, were not associated with elevated β-γ PAC. Finally, medication-induced high β-γ PAC reduction within the supplementary motor area was associated with clinical improvement. Altogether, this study provides novel insights into the pathophysiology of PD as an oscillopathy and identifies high β-γ PAC as a potential cortical marker of Parkinsonian symptoms and treatment effects. This has important implications for invasive as well as non-invasive therapeutic strategies, as high β-γ PAC targeting might hold greater promise than targeting β-γ PAC per se.

### High β-γ PAC is elevated in the cortical motor network

In PD, elevated β-γ phase-amplitude coupling has been detected in cortical and subcortical structures using invasive and non-invasive techniques (Karekal *et al*., 2022). In particular, elevated β-γ PAC was observed within the STN and associated with symptom severity of bradykinesia and rigidity (van Wijk *et al*., 2016). On the cortical level, β-γ PAC across the beta frequency range (13-30 Hz) was elevated over the sensorimotor cortex as assessed by ECoG as well as EEG and was responsive to dopamine replacement therapy and DBS (de Hemptinne *et al*., 2013; de Hemptinne *et al*., 2015; Swann *et al*., 2015). In line with previous findings, we identified elevated β-γ PAC within areas of the cortical motor system. We extend these findings by identifying a frequency-specific elevation of PAC, including high-β to broadband-γ PAC. Low-β-γ PAC, however, was not elevated in PD patients. The β-band has been subdivided into low-(13-22 Hz) and high-β (23-35 Hz) to differentiate the roles of distinct frequencies in PD symptoms and to provide a more nuanced understanding of neural oscillations in normal and pathological states. In particular, Binns and colleagues show that high beta power within the primary motor cortex can be associated with bradykinesia and decreased after dopaminergic treatment. While similar observations have been reported in rodents, it’s important to note that human beta oscillations have distinct frequency characteristics that may not directly translate from rodent findings (Dupre *et al*., 2016; Binns *et al*., 2024). At the subcortical level, STN low-beta oscillations have been associated with UPDRS-III total scores, while high-beta power predicted UPDRS-III response to levodopa and DBS therapy (Morelli and Summers, 2023). Moreover, recent studies employing invasive recordings suggest an association between cortico-subcortical coupling in beta sub-bands and anatomical pathways (Oswal *et al*., 2016; Oswal *et al*., 2021; Binns *et al*., 2024). While M1-STN coupling in the low-beta band is supposed to be a surrogate of indirect pathway activity, M1-STN high-beta coupling might be a surrogate of hyperdirect pathway activity, which has been hypothesised to mediate the therapeutic effects of DBS and dopaminergic therapy (Binns *et al*., 2024). Using invasive recordings, such as electrocorticography, bears the advantage of high spatial resolution and low signal-to-noise ratio. Spatial coverage, however, is limited due to the size of the electrode strip that can be placed through a burr hole, and the widespread employment in clinical practice is limited due to its invasive nature (Gong *et al*., 2021). Using advanced source localisation techniques, we overcame the limitations of spatial coverage and the invasive nature of ECoG recordings. Notably, the present results show that elevated high β-γ PAC is present not only in somatosensory areas but extends to additional areas of the cortical motor system, namely PMC. This is in line with the results by Gong and colleagues, who showed that β-γ PAC was elevated in cortical sources, including PMC and BA3 (Gong *et al*., 2021). Our results corroborate the notion that β-γ PAC is a potential marker for the Parkinsonian state but highlight the importance of subdividing the beta band to allow for more precise approaches in differentiating between healthy controls and PD patients. Moreover, we further extend the findings by Gong and colleagues by showing that β-γ PAC was elevated between sources of the cortical motor network. Functional and induced connectivity analyses showed distinct alterations in PD patients within the cortical motor network (Nettersheim *et al*., 2019; Loehrer *et al*., 2022). In particular, enhanced beta-gamma coupling between primary motor cortices was associated with poor motor performance and prefrontal to premotor coupling is pathologically altered in PD (Nettersheim *et al*., 2019). Accordingly, we found exaggerated high β-γ PAC between prefrontal and premotor areas and between prefrontal cortex and primary motor cortex in PD patients. As effect sizes of post-hoc comparisons of high β-γ PAC between HC and PD patients were larger within cortical sources compared to between sources, we argue that high β-γ PAC within PMC and M1 is a more suitable marker to differentiate between PD patients and healthy controls. In line with previous studies, we found no difference in cortical beta power between healthy controls and PD patients, suggesting that cortical high β-γ PAC is a more reliable marker to differentiate between the healthy and diseased state (Litvak *et al*., 2011; de Hemptinne *et al*., 2013). Altogether, these findings reinforce the importance of delineating the frequency-specific characteristics of β-γ PAC within the cortical motor system and shed light on the potential role of high β-γ PAC as a diagnostic marker.

### Dopaminergic therapy reduces elevated high β-γ PAC in the motor network

Given the excessive increase in cortical β-γ PAC and its supposed role as a marker for PD, several studies have investigated the effects of dopaminergic replacement therapy and DBS on PAC. In particular, de Hemptinne et al. showed that DBS reduces β-γ PAC in M1 and suggested a potential role in mediating the clinical effects of DBS (de Hemptinne *et al*., 2015). Furthermore, Miller and colleagues showed an impact of levodopa on PAC at the sensor level (Miller *et al*., 2019). The present study sheds light on multiple aspects of dopaminergic influence on β-γ PAC at the cortical level, most prominently demonstrating that exaggerated high β-γ PAC is reduced in all sources of the cortical motor network by dopaminergic therapy. This corroborates the results by de Hemptinne and Miller and confirms the widespread influence dopaminergic medication exerts on the cortex. The significant interaction effect between medication and frequency identified in our analysis is particularly important. Our results suggest that levodopa exerts a frequency-specific effect at the cortical level, specifically reducing exaggerated high β-γ PAC, while low β-γ PAC is not significantly influenced. Indeed, our results align with recent findings by Binns et al., who showed that levodopa selectively reduced high beta activity over the primary motor cortex in PD (Binns *et al*., 2024). The role of cortical beta oscillations in differentiating between medication states remains controversial, as some studies have shown a reduction of cortical beta power while others show an increase (Stoffers *et al*., 2007; Melgari *et al*., 2014). Therefore, high β-γ PAC may represent a more reliable marker to differentiate between medication states and might be more promising for treatment surveillance. This is further corroborated as we demonstrate a frequency-specific reduction of high β-γ PAC between cortical sources induced by dopaminergic therapy.

### High β-γ PAC is associated with motor symptoms and clinical improvement

Further support for the importance of delineating the frequency-specific dynamics of β-γ PAC comes from our observation that symptom severity was associated with high β-γ PAC. In line with previous studies, our observed associations reinforce the role of β-γ PAC in impaired motor control and highlight its potential as a marker for disease severity (Miller *et al*., 2019; Gong *et al*., 2021). Our findings extend prior research by demonstrating that this association is frequency-specific, as no associations were observed for low β-γ PAC. Furthermore, we show that this frequency-specific association is present within all sources of the motor network except DLPFC. This is particularly relevant for two reasons. First, identifying the specific areas where PAC is associated with clinical symptoms could help refine diagnostic and treatment strategies as therapies targeting PAC could adapt to the spatial and frequency-specific dynamics. Furthermore, we did not observe any associations of β-γ PAC between sources and clinical symptoms. Therefore, assessing high β-γ PAC within sources of the motor network seems more promising to establish an objective diagnostic measure than evaluating PAC between sources. We further delineate the observed associations by showing that high β-γ PAC is associated with bradykinesia and rigidity subscores. At the same time, tremor scores were not related to β-γ PAC within and between sources of the motor network. Our findings align with the long-held position that distinct symptoms are linked to specific patterns of oscillatory activity: while bradykinesia and rigidity scores are associated with increased oscillatory activity in the beta band, tremor scores are supposed to be related to lower frequency activity (Kühn *et al*., 2006; Kühn *et al*., 2009). Finally, we demonstrate an association between therapy-induced changes in PAC and therapy-induced changes in clinical severity. Specifically, a reduction in high β-γ PAC within the SMA was linked to improvements in overall motor symptoms, with a particular impact on reducing bradykinesia and rigidity.

Several considerations have to be taken into account when interpreting the present finding. First, cortical layer 5 neurons in the SMA project monosynaptically to the STN via the hyperdirect pathway (Bingham *et al*., 2023). Recent studies have highlighted this cortical input to the STN via the hyperdirect pathway as a central contributor to pathological synchrony within the STN and between the cortex and STN (Oswal *et al*., 2016; Oswal *et al*., 2021; Binns *et al*., 2024). Furthermore, Oswal and colleagues demonstrated that high beta activity in the SMA drives STN activity via the hyperdirect pathway (Oswal *et al*., 2021), and this pathway has widely been considered the main target for the therapeutic effects of dopaminergic therapy and DBS. Therefore, we suggest that the SMA’s high β-γ PAC is a marker for exaggerated synchrony in the hyperdirect pathway. Accordingly, we argue that high β-γ PAC within the SMA might be a specific neural target for adaptive treatment approaches, whether invasive (such as DBS or pump-therapy) or non-invasive, enabling personalised therapy. Altogether, we argue that high β-γ PAC within the cortical motor network can serve as a marker to differentiate PD patients from healthy controls, might be suitable for objectively assessing disease states, and might serve as an input for adaptive treatment approaches.

### Limitations

It is essential to acknowledge that this research is not without its limitations. First, analyses were conducted on recordings while patients were at rest. Therefore, our findings do not permit a direct association between elevated β-γ PAC and the clinical phenotype. Nonetheless, the observed associations between specific PAC properties and Parkinsonian motor symptoms and between PAC change and symptom improvements suggest that exaggerated PAC likely represents a marker of the Parkinsonian state. Second, a significant challenge in analysing cross-frequency coupling, such as PAC, is the presence of non-sinusoidal sawtooth-like oscillations, which can produce spurious PAC (Seymour *et al*., 2017). The characteristics of an oscillation can be quantified by determining specific temporal features, including rise- and decay-time and the ratio between them. As we observed no differences in these measures across the three groups, our results are unlikely affected by the oscillations’ non-sinusoidal properties. Third, PD patients who participated in this study had the akinetic-rigid and mixed subtypes. No patients with tremor-dominant subtype were included. This restricts the generalizability of our findings across all Parkinson’s disease phenotypes.

## Conclusion

Our study identifies high β-γ PAC as a potential biomarker for the Parkinsonian state and therapeutic strategies. Within the cortical motor network, high β-γ PAC was found to be elevated and subsequently attenuated by dopaminergic replacement therapy. Furthermore, high β-γ PAC was correlated with bradykinesia and rigidity, whereas the medication-induced reduction of high β-γ PAC in the supplementary motor area was associated with clinical improvement. These findings may offer further insight into the pathophysiology of PD as an oscillopathy by elucidating the spatial and frequency-specific characteristics of cortical β-γ PAC. Ultimately, our study provides a possible foundation for future therapeutic interventions targeting specific cortical oscillatory activity.

## Acknowledgement

The authors thank the participants for their active engagement in this study.

## Contributorship

PAL: study concept and design, data acquisition, data analysis, drafting of the manuscript

SY: data analysis, critical revision of the manuscript

IW: data analysis, critical revision of the manuscript

VS: data acquisition, critical revision of the manuscript

SH: critical revision of the manuscript

AP: data analysis, critical revision of the manuscript

LC: data analysis, critical revision of the manuscript

LW: data analysis, critical revision of the manuscript

GRF: critical revision of the manuscript

DJP: data analysis, critical revision of the manuscript

LT: study concept and design, critical revision of the manuscript

HT: study concept and design, data analysis, drafting of the manuscript

