## supplementary material for "Characterizing the frequency-specific and spatiotemporal dynamics of β-γ Phase-Amplitude Coupling in Parkinson’s disease"

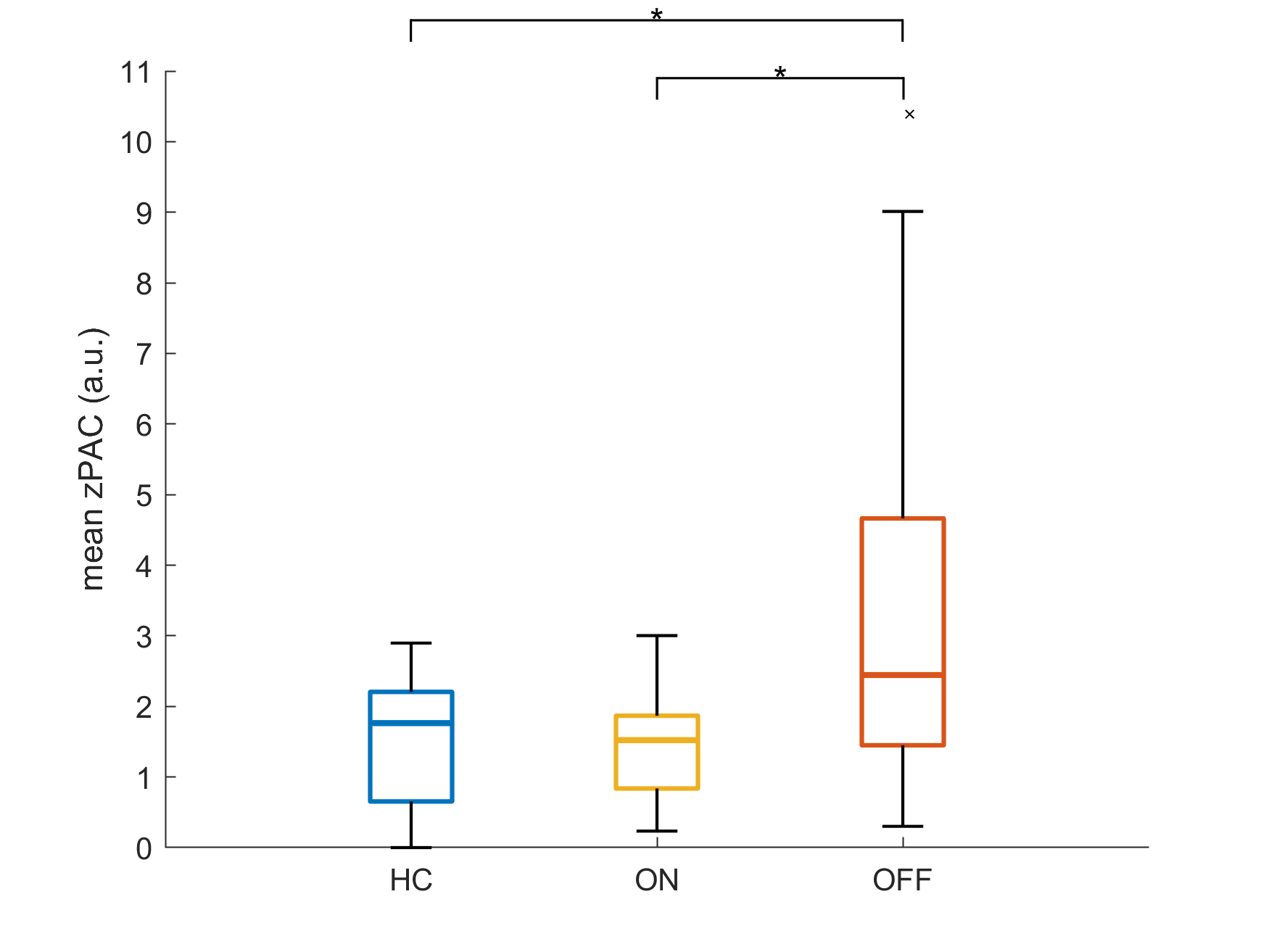

**Supplementary Figure 1.** **Comparison of Mean β-γ PAC across Groups.** Boxplots illustrate the distribution of z-transformed PAC values between the β-frequency range (13-35 Hz) and broadband γ-frequency range (50-150 Hz) across healthy controls (HC), Parkinson’s disease patients in the ON-medication state (PD ON), and those in the OFF-medication state (PD OFF). zPAC values were averaged across hemispheres and regions of interest (M1, lPM, SMA, and DLPFC). The boxes represent the interquartile range (IQR), with the horizontal line indicating the median. Whiskers extend to 1.5 times the IQR or the most extreme data points within this range, while outliers are shown as individual points. **Abbreviations:** a.u. = arbitrary units; HC = healthy controls; PAC = phase-amplitude coupling; OFF = Parkinson’s disease patients in the OFF-medication state; PD ON = Parkinson’s disease patients in the ON-medication state

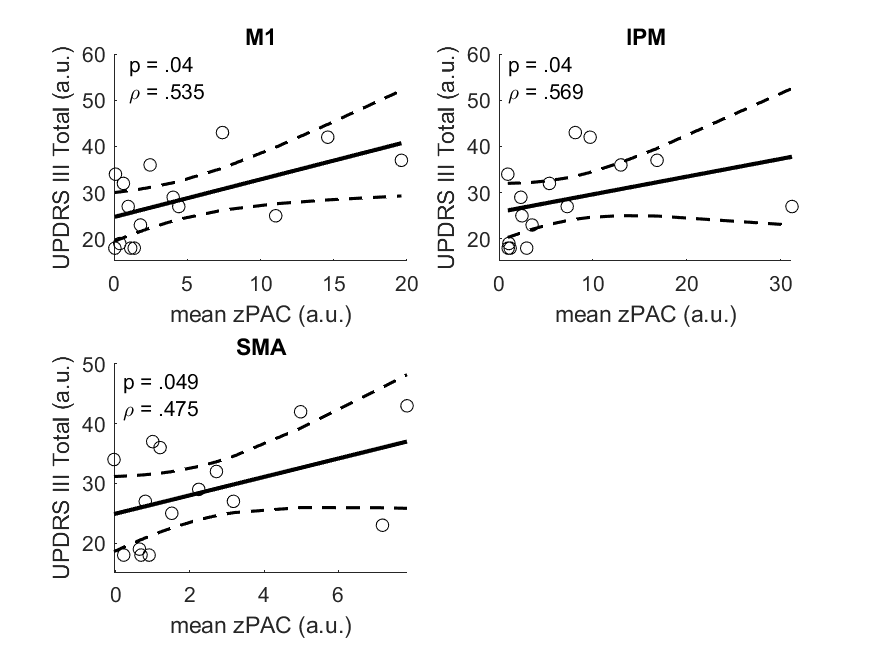

**Supplementary Figure 2.** **Correlation of high β-γ PAC with UPDRS-III Total Scores.** Relationship between high β-γ PAC and UPDRS-III Total scores in the OFF medication state across three regions of interest, with zPAC values averaged across hemispheres. Only regions with significant relationships are presented. Solid black lines represent the best-fit line while dashed lines represent 95% confidence intervals. P-values were corrected for multiple comparisons using the Benjamini-Hochberg method. **Abbreviations:** a.u. = arbitrary units; PAC = phase-amplitude coupling; UPDRS = Unified Parkinson’s Disease Rating Scale

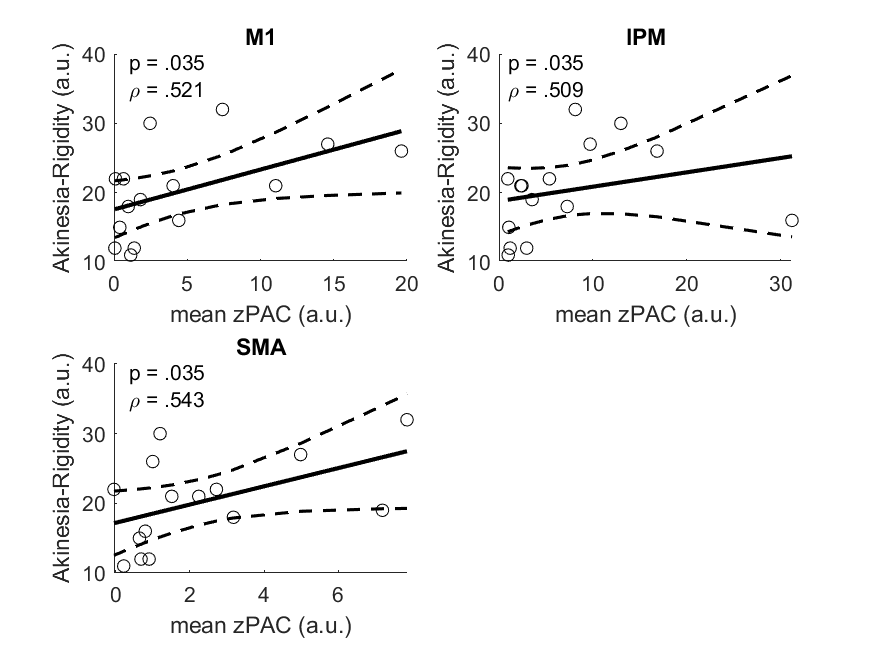

**Supplementary Figure 3.** **Correlation of high β-γ PAC with UPDRS-III Bradykinesia-Rigidity Subscore.** Relationship between high β-γ PAC and UPDRS-III bradykinesia-rigidity subscore in the OFF medication state across three regions of interest, with zPAC values averaged across hemispheres. Only regions with significant relationships are presented. Solid black lines represent the best-fit line while dashed lines represent 95% confidence intervals. P-values were corrected for multiple comparisons using the Benjamini-Hochberg method. **Abbreviations:** a.u. = arbitrary units; PAC = phase-amplitude coupling; UPDRS = Unified Parkinson’s Disease Rating Scale

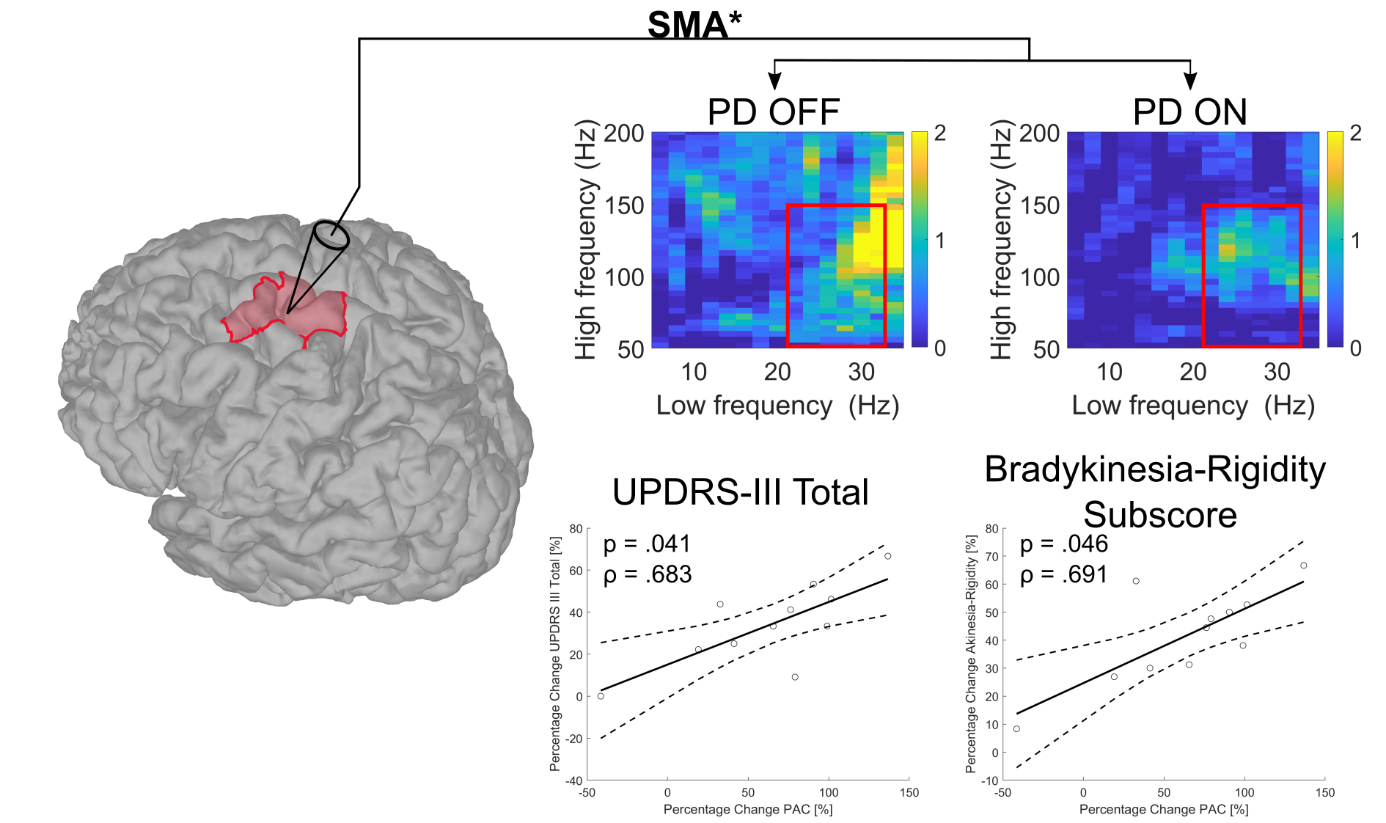

**Supplementary Figure 4.** **Correlation of high β-γ PAC changes in the SMA with UPDRS-III improvement – Analysis without outliers.** Relationship between the percentage change in high β-γ PAC within the SMA of the primarily affected side and the percentage change in UPDRS-III total scores as well as the bradykinesia-rigidity subscore. In this analysis, percentage change PAC outliers were excluded. Greater reduction of high β-γ PAC in SMA was associated with greater reduction in UPDRS-III total scores as well as bradykinesia-rigidity subscores. The solid black line represents the best-fit line while dashed lines represent 95% confidence intervals. P-values were corrected for multiple comparisons using the Benjamini-Hochberg method. **Abbreviations:** PAC = phase-amplitude coupling; OFF = Parkinson’s disease patients in the OFF-medication state; PD ON = Parkinson’s disease patients in the ON-medication state; SMA = supplementary motor area; UPDRS = Unified Parkinson’s Disease Rating Scale

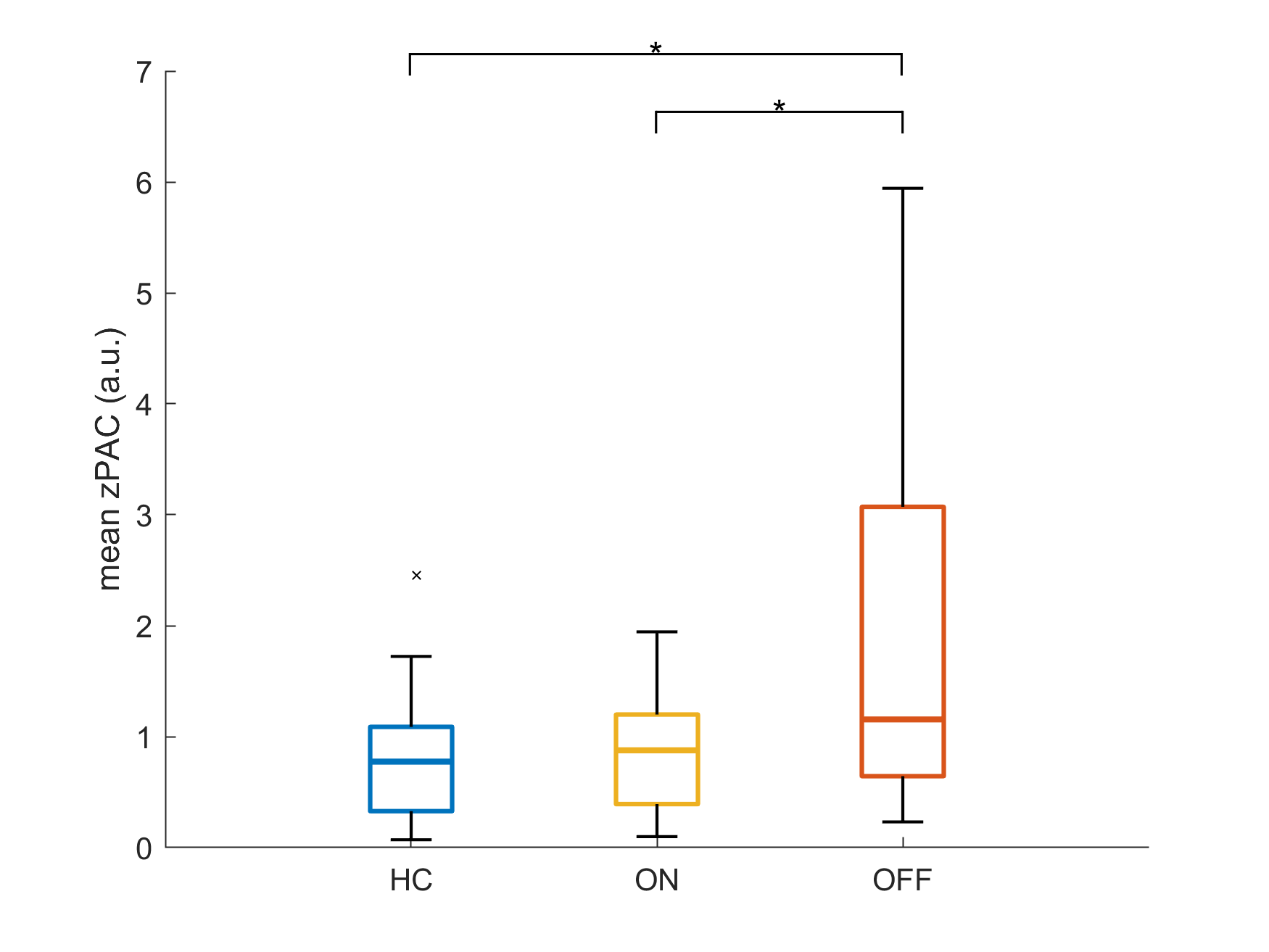

**Supplementary Figure 5.** **Comparison of Mean β-γ PAC between Sources of the Human Motor Network across Groups.** Boxplots illustrate the distribution of z-transformed PAC values between the β-frequency range (13-35 Hz) and broadband γ-frequency range (50-150 Hz) across healthy controls (HC), Parkinson’s disease patients in the ON-medication state (PD ON), and those in the OFF-medication state (PD OFF). zPAC values were averaged across hemispheres and regions of interest. The boxes represent the interquartile range (IQR), with the horizontal line indicating the median. Whiskers extend to 1.5 times the IQR or the most extreme data points within this range, while outliers are shown as individual points. **Abbreviations:** a.u. = arbitrary units; HC = healthy controls; PAC = phase-amplitude coupling; OFF = Parkinson’s disease patients in the OFF-medication state; PD ON = Parkinson’s disease patients in the ON-medication state

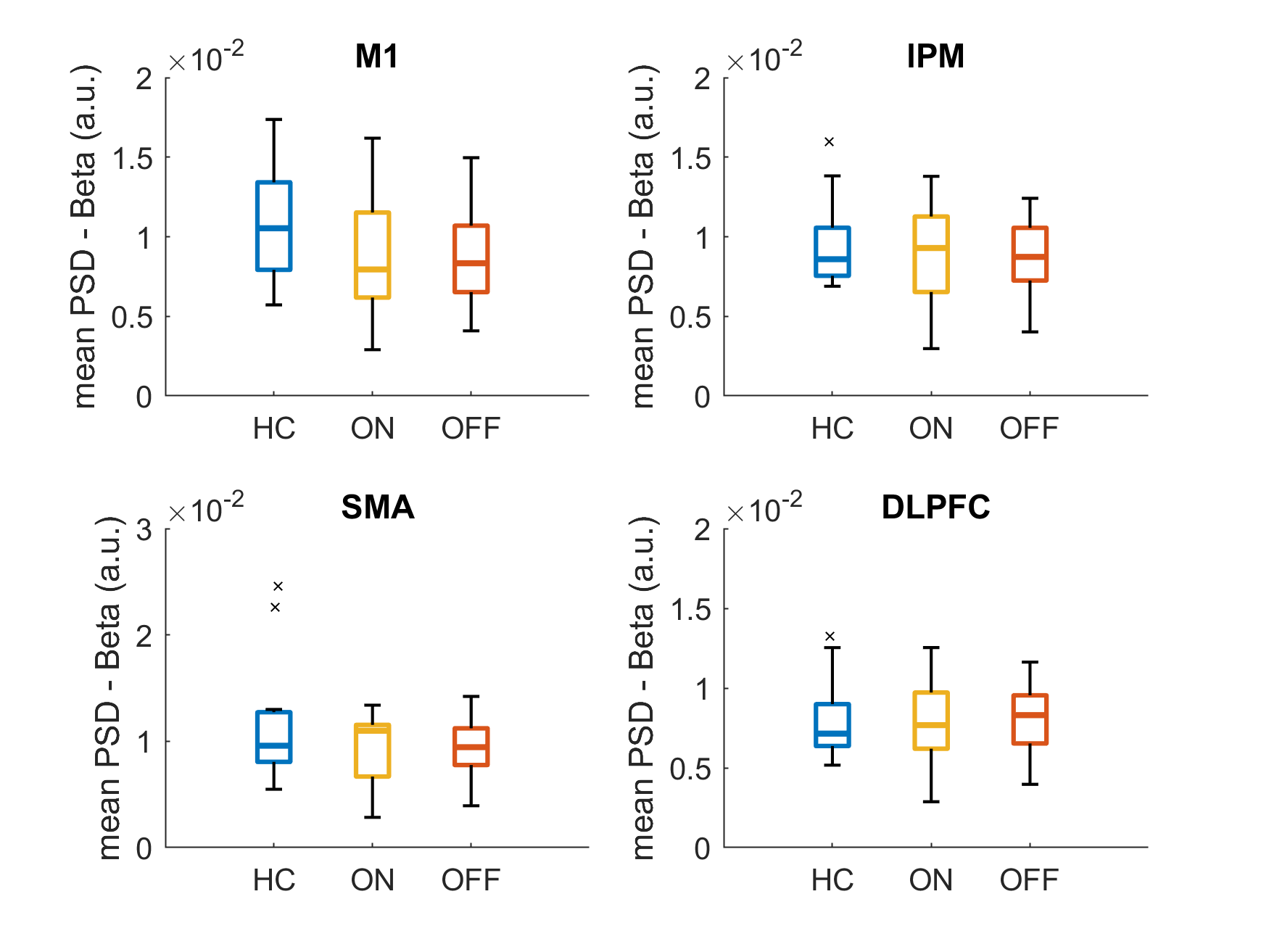

**Supplementary Figure 6.** **Comparison of Mean Beta PSD across Groups.** Boxplots illustrate the distribution of power spectral density (PSD) values in the β-frequency range (13-35 Hz) across healthy controls (HC), Parkinson’s disease patients in the ON-medication state (PD ON), and those in the OFF-medication state (PD OFF). The boxes represent the interquartile range (IQR), with the horizontal line indicating the median. Whiskers extend to 1.5 times the IQR or the most extreme data points within this range, while outliers are shown as individual points. **Abbreviations:** a.u. = arbitrary units; HC = healthy controls; PSD = power spectral density; OFF = Parkinson’s disease patients in the OFF-medication state; PD ON = Parkinson’s disease patients in the ON-medication state;

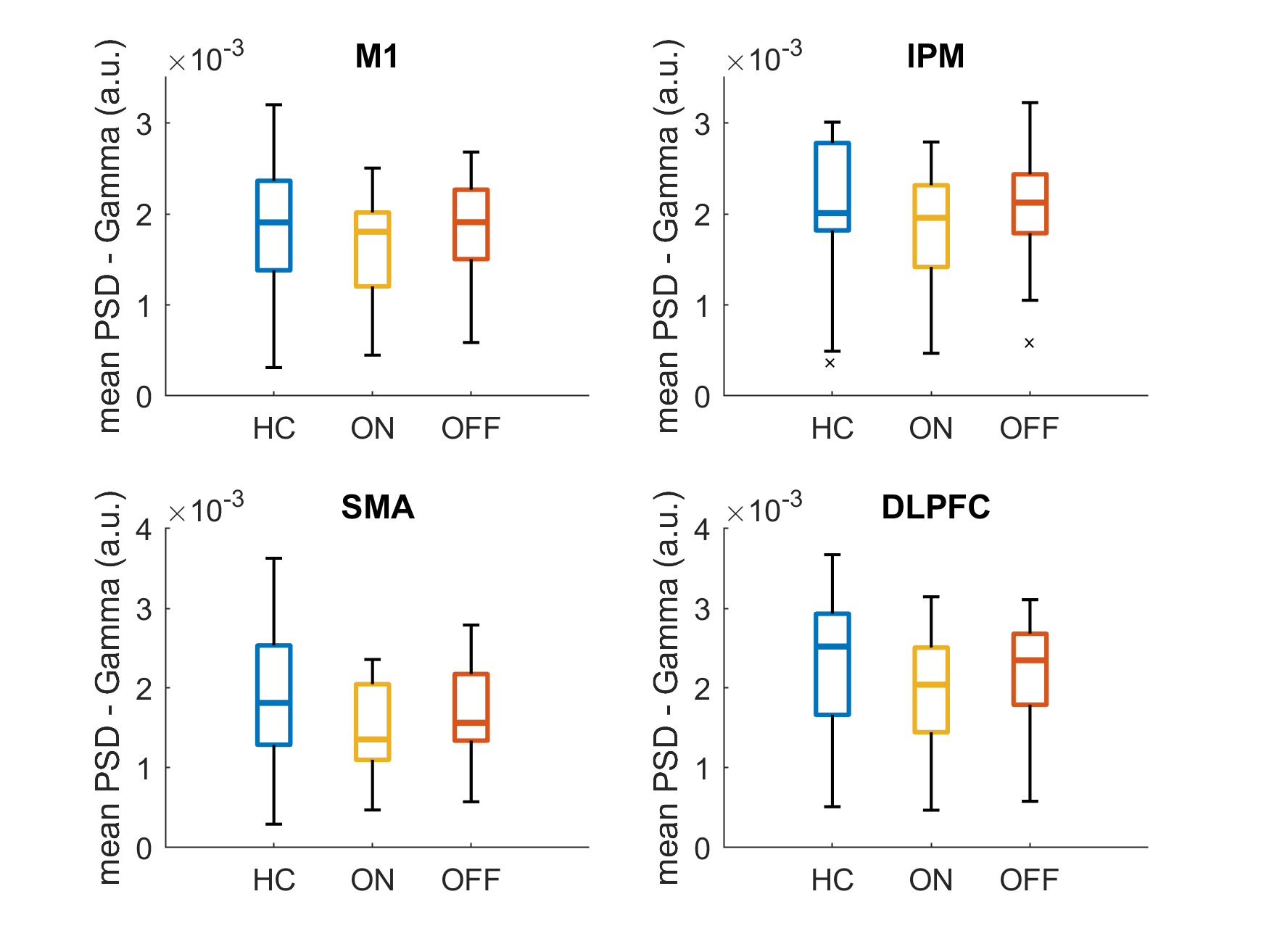

**Supplementary Figure 7.** **Comparison of Mean Gamma PSD across Groups.** Boxplots illustrate the distribution of power spectral density (PSD) values in the γ-frequency range (50-150 Hz) across healthy controls (HC), Parkinson’s disease patients in the ON-medication state (PD ON), and those in the OFF-medication state (PD OFF). The boxes represent the interquartile range (IQR), with the horizontal line indicating the median. Whiskers extend to 1.5 times the IQR or the most extreme data points within this range, while outliers are shown as individual points. **Abbreviations:** a.u. = arbitrary units; HC = healthy controls; PSD = power spectral density; OFF = Parkinson’s disease patients in the OFF-medication state; PD ON = Parkinson’s disease patients in the ON-medication state;

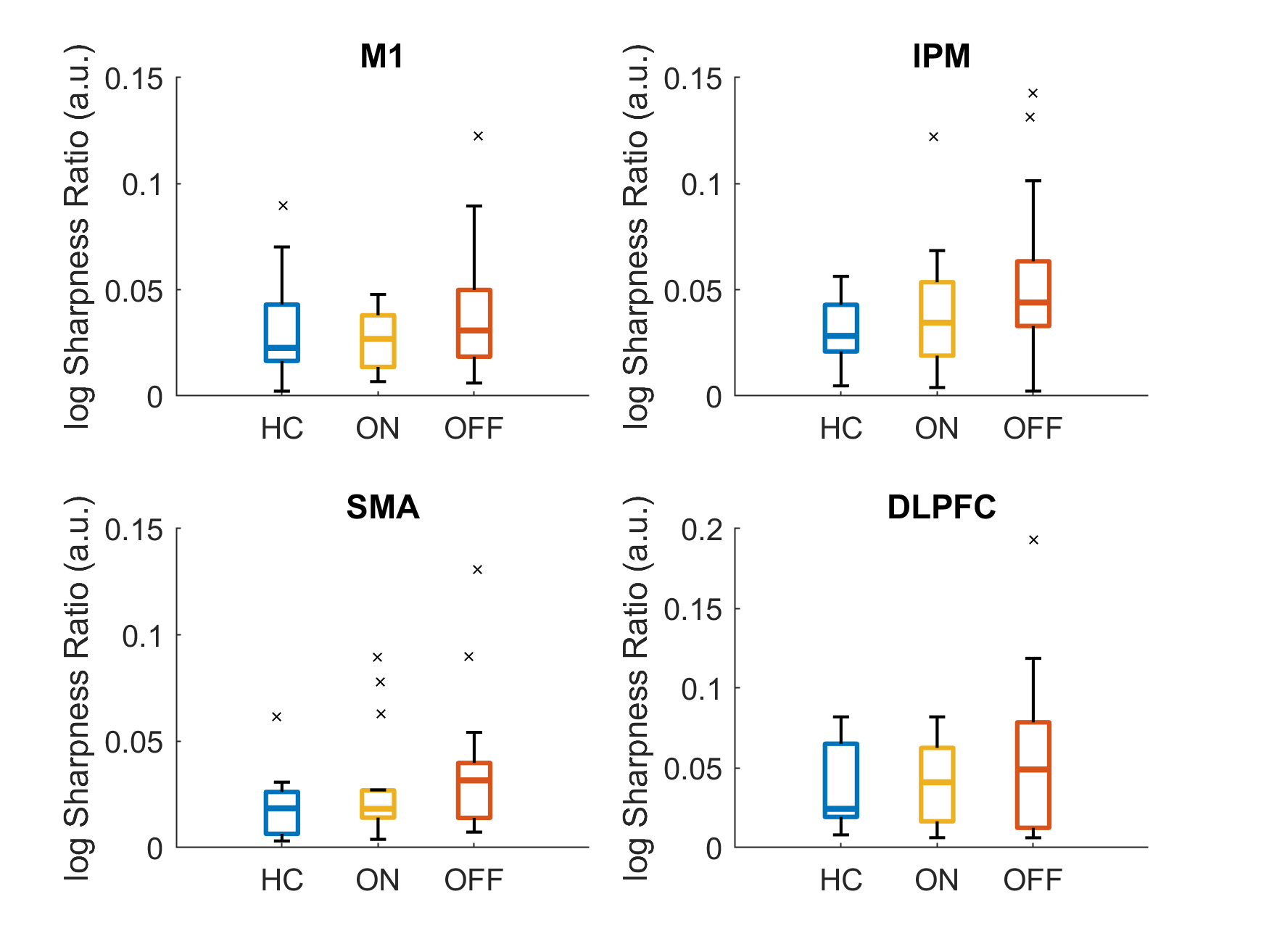

**Supplementary Figure 8.** **Comparison of Sharpness Ratio across Groups.** Boxplots illustrate the distribution of sharpness ratio in the β-frequency range (13-35 Hz) across healthy controls (HC), Parkinson’s disease patients in the ON-medication state (PD ON), and those in the OFF-medication state (PD OFF). The boxes represent the interquartile range (IQR), with the horizontal line indicating the median. Whiskers extend to 1.5 times the IQR or the most extreme data points within this range, while outliers are shown as individual points. **Abbreviations:** a.u. = arbitrary units; HC = healthy controls; OFF = Parkinson’s disease patients in the OFF-medication state; PD ON = Parkinson’s disease patients in the ON-medication state;

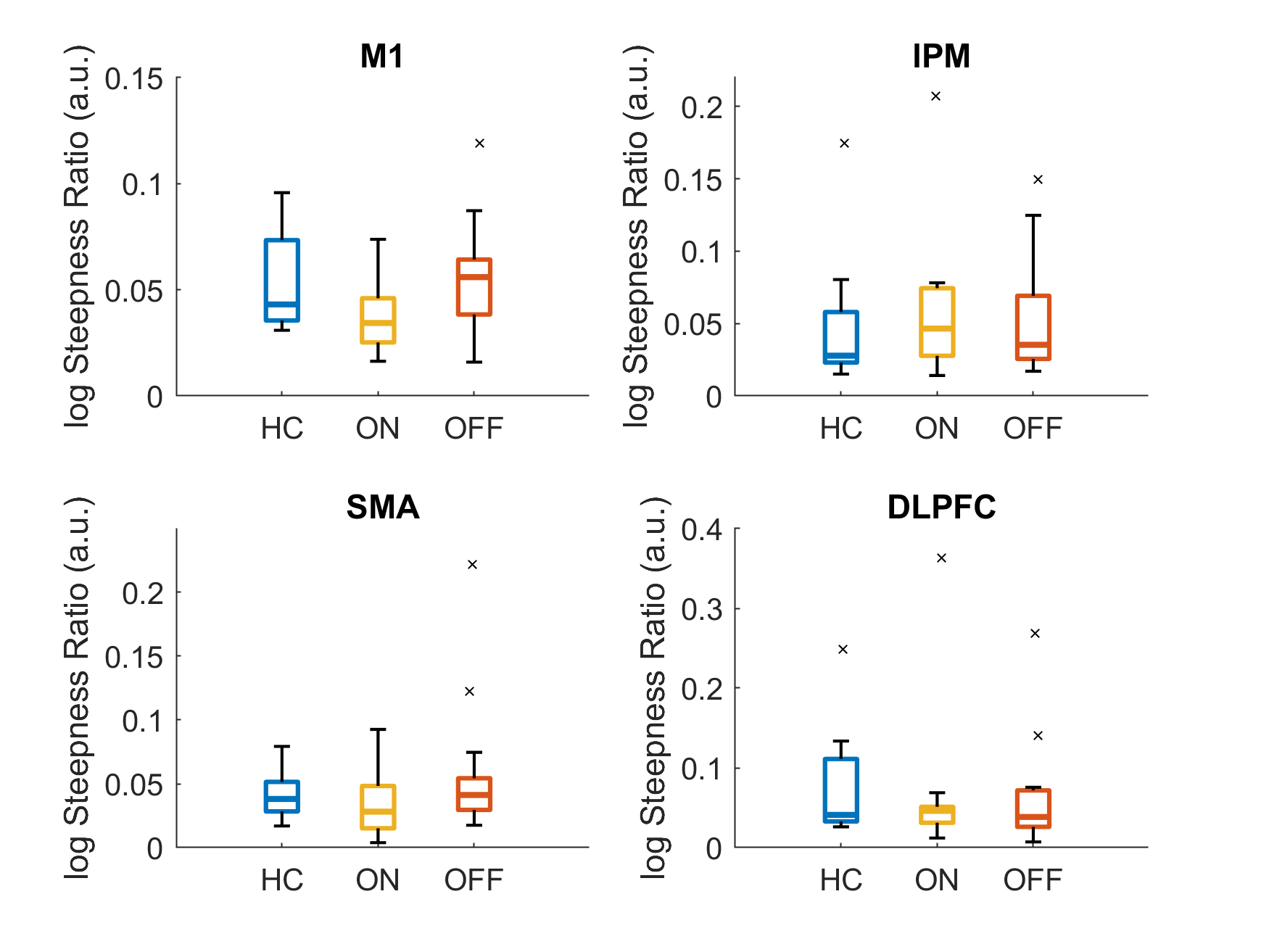

**Supplementary Figure 9.** **Comparison of Steepness Ratio across Groups.** Boxplots illustrate the distribution of steepness ratio in the β-frequency range (13-35 Hz) across healthy controls (HC), Parkinson’s disease patients in the ON-medication state (PD ON), and those in the OFF-medication state (PD OFF). The boxes represent the interquartile range (IQR), with the horizontal line indicating the median. Whiskers extend to 1.5 times the IQR or the most extreme data points within this range, while outliers are shown as individual points. **Abbreviations:** a.u. = arbitrary units; HC = healthy controls; OFF = Parkinson’s disease patients in the OFF-medication state; PD ON = Parkinson’s disease patients in the ON-medication state;

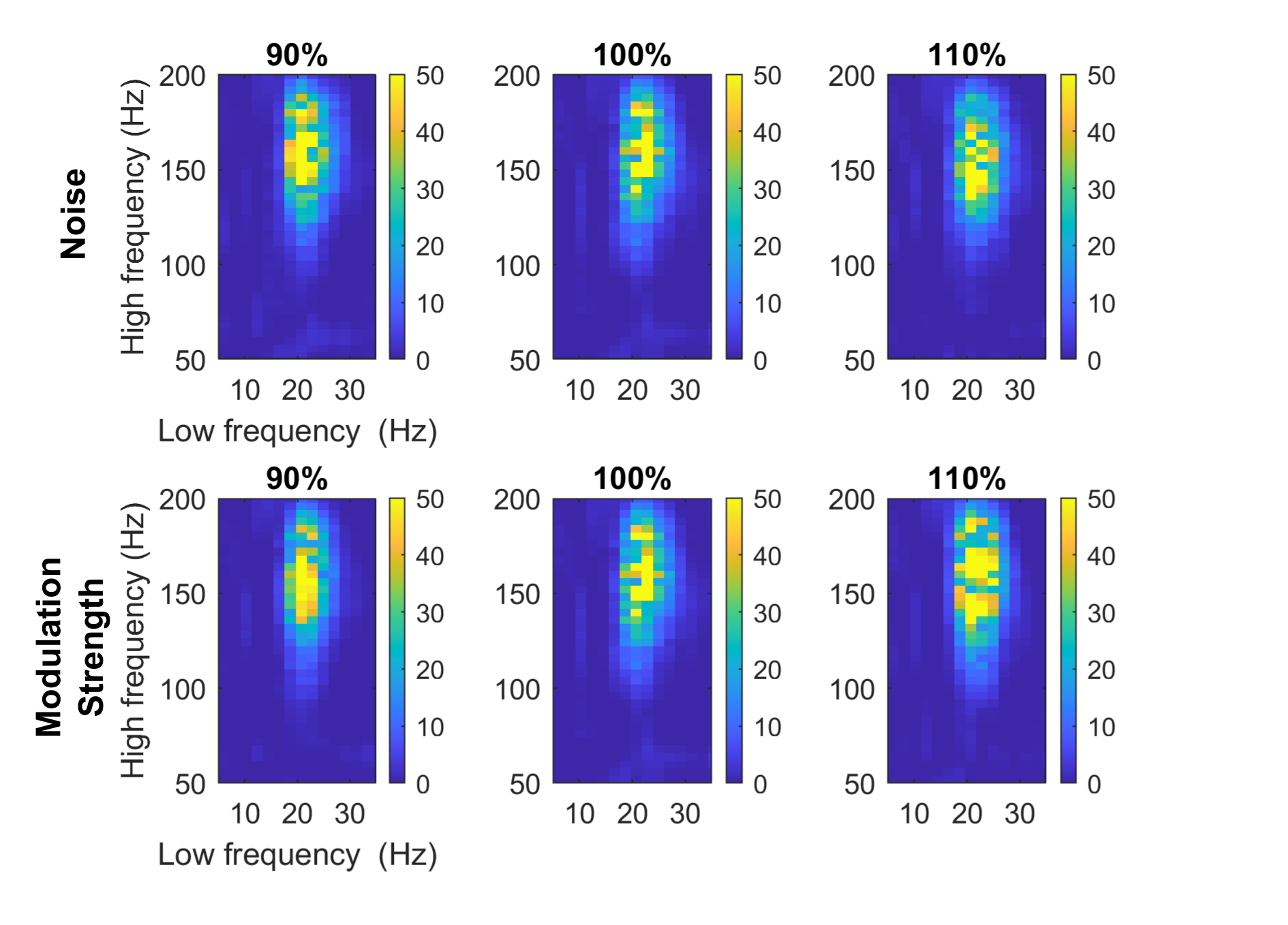

**Supplementary Figure 10.** **Validation of Phase-Amplitude Coupling Detection with Simulated Data.** Simulated data segments of three seconds were generated with known beta-gamma phase-amplitude coupling (21 Hz phase, 100 Hz amplitude center frequencies) at varying levels of random noise and coupling strengths. The same MATLAB analysis pipeline used for EEG data was applied, successfully detecting beta-gamma PAC.
